# Step-wise epigenetic signal integration governs adaptive programming of cytotoxic lymphocytes

**DOI:** 10.1101/2023.11.07.565992

**Authors:** Simon Grassmann, Endi K. Santosa, Hyunu Kim, Isabelle B. Johnson, Adriana M. Mujal, Jean-Benoit LeLuduec, Sherry X. Fan, Mark Owyong, Adrian Straub, Liang Deng, Katharine C. Hsu, Dirk H. Busch, Colleen M. Lau, Joseph C. Sun

## Abstract

Lymphocyte differentiation depends on activation via antigen and cytokines during the immune response to infection. How the timing and integration of these signals program the epigenetic and functional fate of these cells is not completely understood. In this study, we find that interleukin (IL)-12 signaling received by innate and adaptive lymphocytes has a context-dependent role for immune memory formation. In the absence of a preceding and/or sufficient antigen receptor signaling event, IL-12 impairs the adaptive expansion of cytotoxic lymphocytes. In contrast, sufficient antigen-receptor signaling redirects inflammatory cytokine signals to promote memory differentiation via cooperation of STAT4 and AP-1 transcription factors. By this crucial epigenetic mechanism, optimally equipped lymphocytes are selected for memory formation rather than a terminal effector cell fate. Whereas T cells are hardwired to be shielded from premature IL-12 signaling, NK cells rely on coincidental early antigen receptor signaling for adaptive responses. Together, step-wise integration of antigen and cytokine signaling optimizes both effector and memory differentiation, allowing for promiscuous recruitment into the acute immune response while promoting avidity maturation in memory populations of both innate and adaptive lymphocytes.

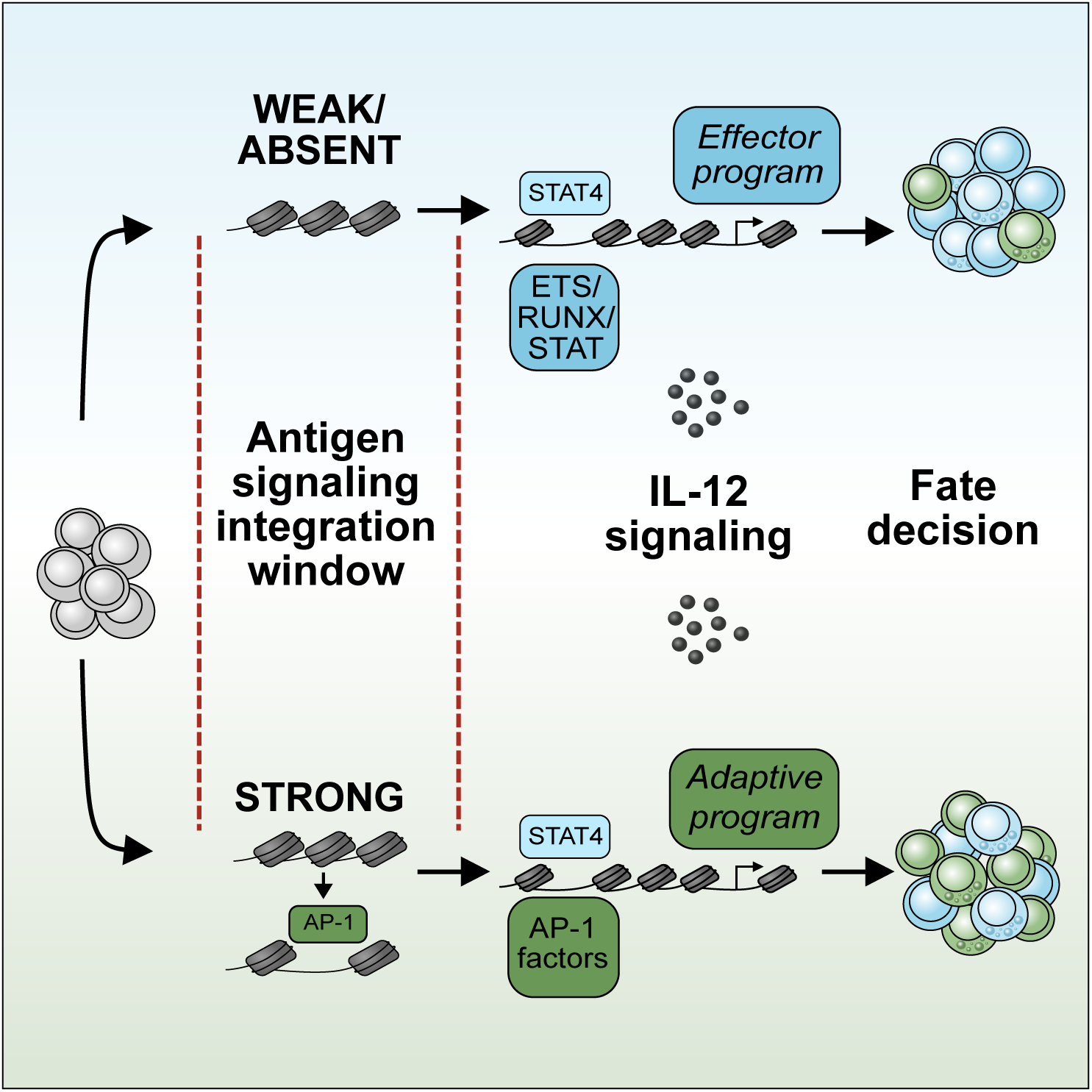

**Key points:**

- Adaptive NK cell responses rely on sequential integration of antigen and inflammatory signals.
- Epigenetic redirection of STAT4 genomic binding promotes adaptive programming.
- CD8^+^ T cell fate depends on antigen-dependent integration of inflammatory signaling.
- STAT/AP-1 cooperation underlies step-wise integration of antigen and cytokine signaling in NK cells and CD8^+^ T cells.

## Introduction

Adaptive immune responses are crucial for protection against infections and cancer. During infection, immune cells must mount a robust effector response while also creating long-lived memory cells that protect against re-infection. Adaptive lymphocyte responses rely both on activating receptor and cytokine signaling^1,2^. How these signals are integrated to optimally tailor effector versus memory differentiation is incompletely understood. In T cells, antigen signaling and concomitant inflammatory cytokine signaling shapes effector differentiation into memory precursors and short-lived effector cells^3,4^. Strong antigen-receptor signals promote both short-lived effector and memory differentiation to drive avidity maturation of memory T cell populations^5^. In contrast, inflammatory cytokines promote differentiation of short-lived effector cells^6^; however, their impact on memory precursor formation remains controversial. Inflammatory cytokine signals have been shown to either reduce^7^, promote^8^ or have no impact^9^ on immunological memory in T cells, suggesting that other factors may influence how inflammatory cytokines affect T cell differentiation.

Recent findings have demonstrated that adaptive immune responses are not restricted to T and B cells of the adaptive immune system. Innate lymphocytes such as NK cells can mount robust adaptive responses during cytomegalovirus (CMV) infection in mice and humans^10,11^. Similar to T cells, adaptive NK cells rely on antigen-receptor signaling^12,13^, show antigen-dependent avidity maturation^14,15^, and require concomitant cytokine signals^16–18^ for their clonal expansion and memory formation. Although NK cells express a variety of activating receptors^19,20^, CMV infection is the main driver of such adaptive responses in NK cells. This suggests that CMV infection provides either a unique combination or sequence of signals leading to adaptive responses by otherwise innate lymphocytes. Originally, it was proposed that NK cells during MCMV infection first participate in a cytokine-driven innate response before mounting their adaptive responses^12^. However, the exact sequence of antigen and cytokine signals has not been tested *in vivo*. We hypothesize that understanding the molecular events underlying adaptive NK cell responses during CMV infection can elucidate the core principles of lymphocyte adaptive responses. In our study, we find that adaptive NK cell responses rely on sequential and epigenetic integration of antigen and STAT4 signaling. By the same mechanism, STAT4 shapes the fate decisions of high and low avidity CD8^+^ T cells.

## Results

### Antigen receptor signal precedes cytokine signals in adaptive NK cells

Because proinflammatory cytokine signals are crucial for lymphocyte responses in addition to antigen receptor signals^16–18^, we hypothesized that one of two mechanisms mediates signal integration (**Supp. Fig. 1A**): 1) Priming of lymphocytes involves concomitant antigen receptor and cytokine signaling (<u>simultaneous integration</u>) or 2) Antigen receptor and cytokine signaling occur sequentially (<u>sequential integration</u>). To determine how antigen and cytokine signals are integrated during adaptive NK cell responses, we analyzed the precise sequence of signaling events in NK cells during mouse cytomegalovirus (MCMV) infection using scRNA-seq (**Fig. 1A** and **Supp. Fig. 1B**). The time course revealed a cluster containing the earliest virus-specific transcriptomic changes at day 1 post-infection (PI) compared to uninfected mice (**Fig. 1B and Supp. Fig. 1B**). Using a cytokine signaling score^21^ versus an antigen-receptor signaling score (**Supp Fig. 1C-D**), we observed that the earliest transcriptomic changes found on day 1 PI were best characterized by antigen-receptor signaling. A robust expression of cytokine-dependent genes did not appear until day 2 PI. Together, the data suggested that antigen receptor signals precede cytokine signals in a subset of NK cells during MCMV infection. The earliest detectable transcriptomic changes included the gene encoding for the antigen receptor Ly49H and genes induced by antigen receptor signaling, such as Nr4a and Egr-family members (**Fig. 1C** and **Supp. Fig. 2A**).

**Figure 1:**
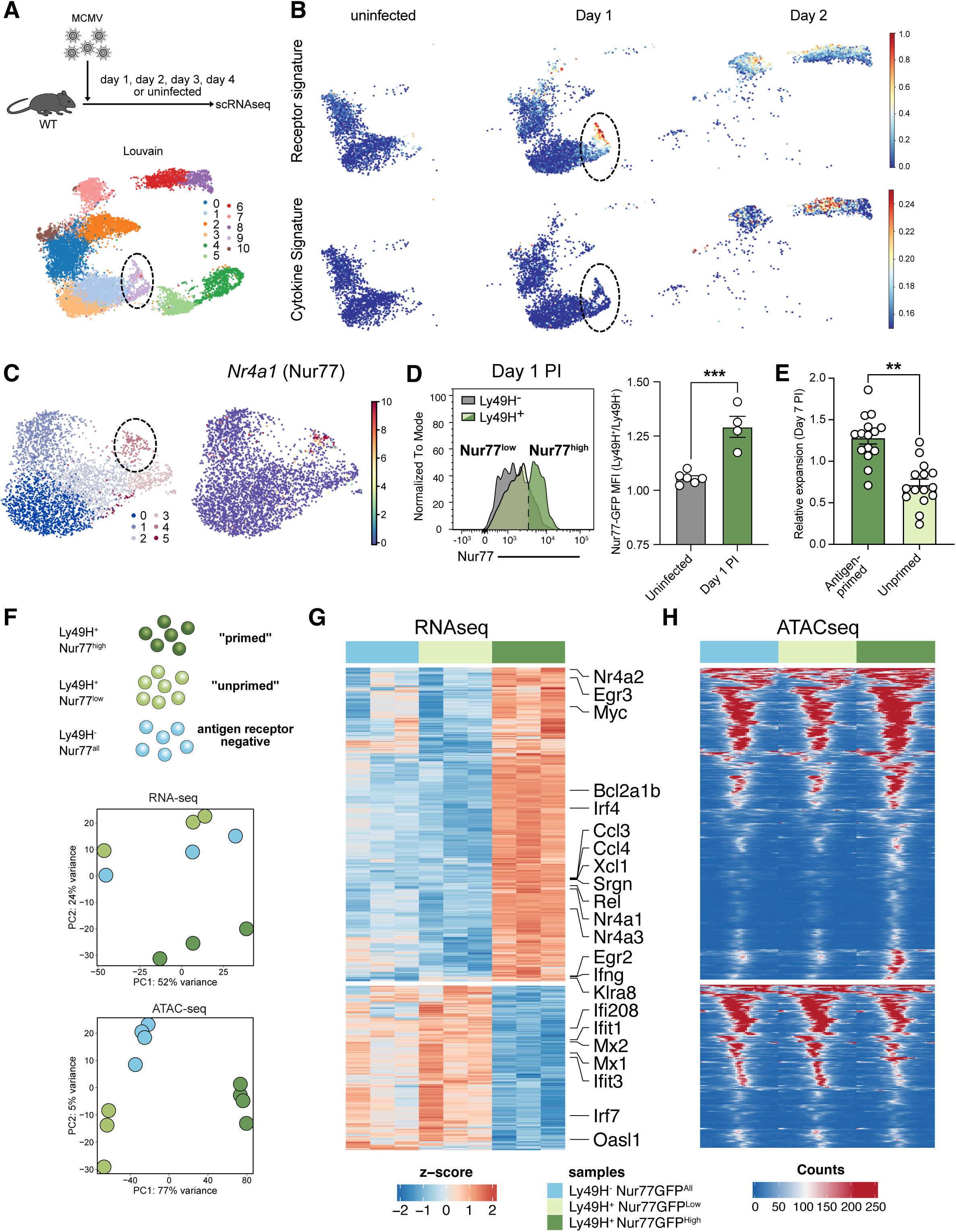
Adaptive NK cell responses rely on early antigen sensing *in vivo*. **A)** UMAP projection of single-cell transcriptomes of total NK cells (Ly49H^+^ and Ly49H^-^) and Louvain Clustering of scRNA-seq analysis and analysis of hashtag proportions in each identified cluster. Dotted line indicates cluster 9, earliest transcriptomic changes on day 1. **B)** Cytokine and antigen receptor score in scRNA-seq separated according to day PI. **C)** Left: Isolated Louvain clustering of single-cell transcriptomes from day 1 PI. Dotted line indicates cluster 4, corresponding to cluster 9 in the full data set. Right: Expression of Nr4a1 in UMAP projection. **D)** Nur77-GFP signal in uninfected mice and on day 1 PI. MFI ratios of Nur77-GFP in Ly49H^+^ and Ly49H^-^ NK cells on day 1 PI. Quantification of Nur77GFP MFI in antigen receptor positive and negative NK cells (ratio Ly49H^+^/Ly49H^-^) **E)** Co-transfer of congenically marked Ly49H^+^ Nur77-GFP^high^ and Ly49H^+^ Nur77-GFP^low^ NK cells into infection-matched Ly49H^-/-^ mice and measurement of expansion. Relative expansion is calculated by dividing % at time of analysis by % at time of injection for comparison between multiple experiments (e.g. 75% at day 7 PI and 50% at time of injection: 75%/50% = 1.5). **F)** Sort of Ly49H^+^ Nur77^high^, Ly49H^+^ Nur77^low^ and Ly49D^+^ Ly49H^-^ NK cells on day 1 PI. PCA plots of RNA-seq and ATAC-seq from Ly49H^+^ Nur77^high^, Ly49H^+^ Nur77^low^ and Ly49D^+^ Ly49H^-^ NK cells on day 1 PI. **G)** Heatmap of RNA-seq of Ly49H^+^ Nur77^high^, Ly49H^+^ Nur77^low^ and Ly49D^+^ Ly49H^-^ NK cells. **H)** Heatmap of ATAC-seq of Ly49H^+^ Nur77^high^, Ly49H^+^ Nur77^low^ and Ly49D^+^ Ly49H^-^ NK cells. Data in **1D)** – **1E)** are pooled from two individual experiments. Error bars represent SEM. Significances are calculated as unpaired or paired T-test. **** p < 0.0001, *** p < 0.001, ** p < 0.01, * p < 0.05.

Because *Nr4a1*, a well-known target of antigen receptor signaling^22^ (**Fig. 1C**), was upregulated on day 1 PI, we used the Nr4a1 (Nur77) GFP reporter mouse^23^ to study the fate of early antigen-primed NK cells. Under steady state, Ly49H^-^ and Ly49H^+^ NK cells showed similar levels of Nur77 (**Supp. Fig. 2B**). In accordance with the scRNA-seq kinetics, Ly49H^+^ NK cells induced Nur77-GFP as early as day 1 PI, with GFP expression enriched in antigen-receptor positive (Ly49H^+^) NK cells compared to antigen-receptor negative (Ly49H^-^) NK cells (**Fig. 1D**). Nur77-reporter expression and scRNA-seq revealed that not all Ly49H^+^ NK cells received this early antigen receptor signal, and suggested there are antigen-receptor primed and unprimed NK cells within the Ly49H^+^ population on day 1 PI. To test whether these two groups differed in their capacity for clonal expansion, we co-transferred equal numbers of Nur77^high^ and Nur77^low^ NK cells at day 1 PI into infection-matched mice and measured their relative numbers 7 days later. In this direct comparison, Nur77^high^ NK cells showed significantly greater expansion than Nur77^low^ NK cells (**Fig. 1E**). Thus, the population of early antigen-primed NK cells represented the main source of the adaptive NK cell pool.

Next, we performed RNA-seq and ATAC-seq on 3 subsets of NK cells on day 1 PI: Ly49H^-^ (antigen receptor negative), Ly49H^+^ Nur77^high^ (antigen primed), and Ly49H^+^ Nur77^low^ (unprimed) NK cells (**Fig. 1F**). Ly49H^+^ Nur77^high^ NK cells showed a transcriptional and epigenetic profile distinct from both Ly49H^+^ Nur77^low^ and Ly49H^-^ NK cells (**Fig. 1G** and **1H**), which were remarkably similar. Furthermore, this Nur77^high^ profile strongly mapped to the observed transcriptomic changes by scRNA-seq on day 1 PI, and was enriched in a distinct cluster on day 2 PI, likely representing antigen-primed NK cells at this timepoint (**Supp. Fig. 2C and 2D**). Ly49H^+^ Nur77^high^ NK cells showed enrichment of antigen-dependent genes and pathways, but reduced expression of genes from cytokine pathways (**Supp. Fig. 3A**). Moreover, comparison of transcriptomic and epigenetic changes in Ly49H^+^ Nur77^high^ NK cells with *in vitro* stimulated NK cells suggested a high degree of correlation only with antigen-receptor signaling (**Supp. Fig. 3B-E**). Together, our data favors the sequential signal integration model, where antigen receptor engagement precedes cytokine signaling to drive adaptive NK cell responses.

### Antigen receptor signals redirect STAT4 genomic binding towards AP-1 sites

To assess whether antigen receptor signaling alters genomic sites bound by cytokine-dependent STAT transcription factors, we compared differentially accessible regions in early antigen primed NK cells to STAT1, STAT4 and STAT5 ChIP-seq datasets from cytokine-stimulated NK cells^21^. Nur77^high^ NK cells showed an enrichment of regions that were bound by STAT4 and, to a lesser degree STAT5 (**Fig. 2A**). To directly test whether antigen stimulation redirects STAT1, STAT4, and STAT5, we sequentially stimulated NK cells with antigen followed by specific STAT-activating cytokines (i.e. IFN-α for STAT1, IL-12 for STAT4, and IL-15 for STAT5) and performed CUT&RUN analysis (**Fig. 2B**). For antigen stimulation, we selected anti-NK1.1 because signaling through NK1.1 and Ly49H elicits similar transcriptomic and epigenetic changes in NK cells (**Supp Fig. 3F**), but anti-Ly49H would only trigger half of the NK cells. Of the three STATs, STAT4 showed the most pronounced increase in genomic binding (**Fig. 2C**), confirming our *in silico* prediction.

**Figure 2:**
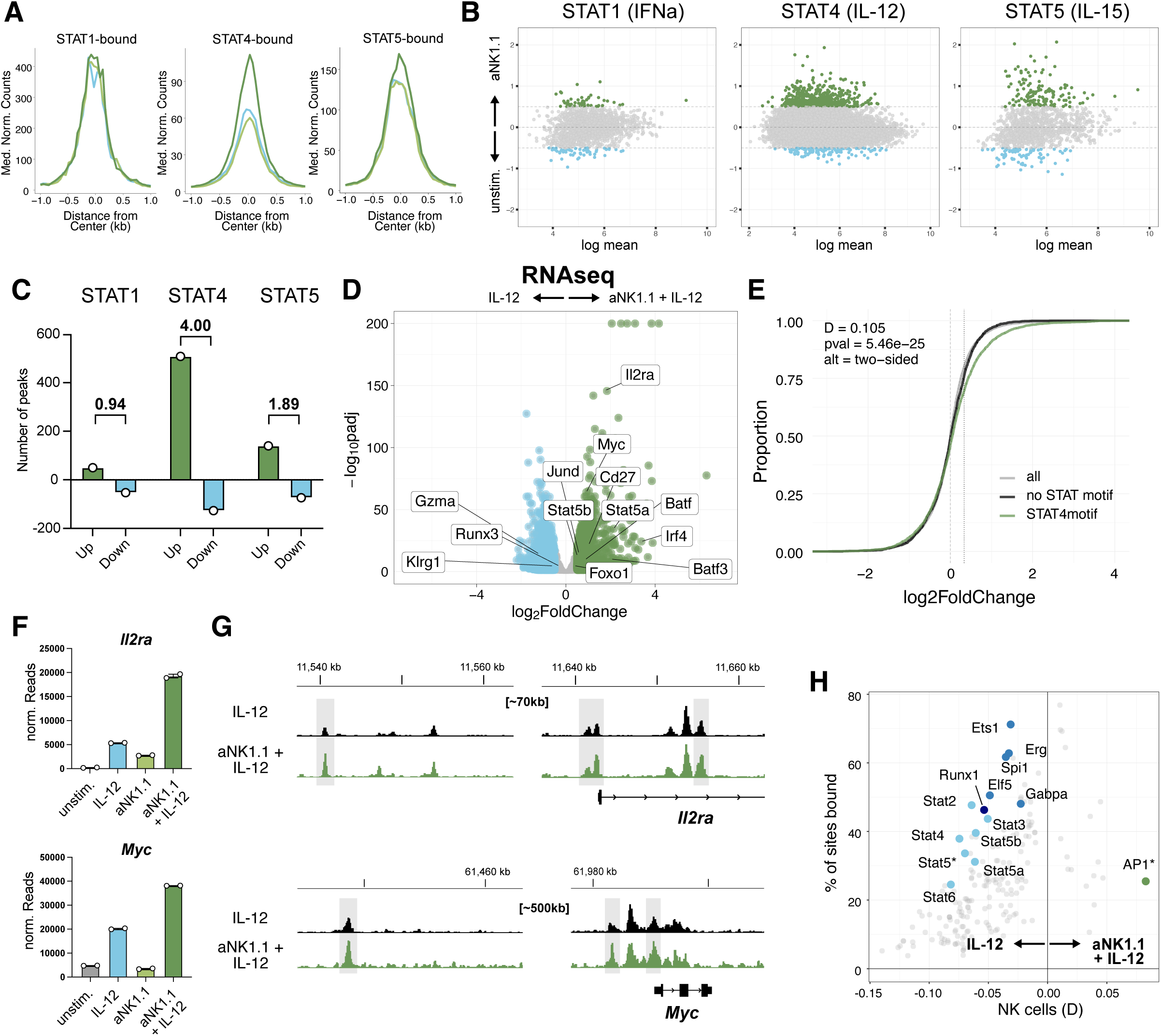
Antigen priming modulates subsequent cytokine signaling by altering STAT4 genomic binding. **A)** Comparison of accessibility of STAT1, STAT4 and STAT5 binding peaks as assessed with ChIPseq. Dark green: Ly49H^+^ Nur77^high^. Light green: Ly49H^+^ Nur77^low^. Light Blue: Ly49H^-^. **B)** NK cells were pre-stimulated with antigen or left untreated. After 3h, NK cells were stimulated with IFN-a, IL-12, or IL-15, and STAT1, STAT4, or STAT5 CUT&RUN, respectively, were performed and analyzed with median-of-ratios normalization. MA plots show mean read count (x-axis) and log2foldchange between the two conditions (y-axis). Peaks with a log2foldchange >0.5 were highlighted. **C)** Quantification of number of peaks with log2foldchange>0.5 in B). Numbers indicate ratios of the peak number that showed positive vs. negative log2foldchange. **D) (left)** Schematic of sequential stimulation with antigen-receptor signaling and IL-12 for RNAseq analysis. **(right)** Volcano plot of differential gene expression in IL-12 stimulated and aNK1.1+IL-12 stimulated NK cells. **E)** ECDF plot of genes that show differential STAT4 binding. Statistics are derived from Kolmogorov-Smirnov test. Genes that showed antigen-dependent STAT4 binding were subsetted into regions that contained a STAT motif or not to discern direct from indirect STAT4 genomic binding. Genes that showed antigen-dependent STAT4 binding and contained a STAT motif showed increased expression in RNA-seq (IL-12 versus aNK1.1 + IL-12). **F)** Example reads for *Il2ra* and *Myc*. **G)** Example normalized STAT4 CUT&RUN tracks averaged across all replicates for *IL2ra* and *Myc* loci in IL-12 stimulated (black) and aNK1.1+IL-12 stimulated (green) NK cells. **H)** Motif enrichment plot. Peaks with antigen-dependent increase and decrease in STAT4 binding in IL-12 and aNK1.1+IL-12 stimulated NK cells were compared with the total STAT4 atlas to examine motif enrichment using one-sided Kolmogorov-Smirnov (KS) test. The KS test statistic D is shown on the x axis and the proportion of regions associated with the motif is shown on the y axis. The odds ratio (frequency of the motif in increase or decrease group divided by its frequency in the entire atlas) was used to assign the motif to either left (regions with antigen-dependent decrease, negative value), or right (regions with antigen-dependent decrease, positive value) in x axis. All motifs (224) from JASPAR CORE(v2024) *mus musculus* database were analyzed. Each color represents motif families of interest as mentioned (STAT, RUNX and ETS motifs). *AP-1 motif labelled here overlaps three motifs, Atf3, Jun and Fos::Jun.

The observed relocation of STAT4 following prior antigen stimulation suggested that antigen signaling altered subsequent IL-12 signaling. To match genomic STAT4 binding to gene expression, we stimulated antigen-receptor pre-stimulated or control NK cells with IL-12 and performed RNA-seq (**Fig. 2D and Supp. Fig. 4A and 4B**). Genes bound by STAT4 showed altered gene expression dependent on direct STAT4 binding, as genes with STAT4 binding that did not harbor a STAT motif (likely via indirect binding) did not show such regulation (**Fig. 2E**). Among the top genes differentially expressed between IL-12-stimulation alone versus sequentially-stimulated NK cells, we found several genes crucial for adaptive NK cell responses, including *Myc*^24^, *Irf4*^25^*, Stat5a, Stat5b* and the high-affinity IL-2 receptor *IL2ra*^16^(**Fig. 2D**). For many of these genes, sequential stimulation increased expression beyond what would be expected if the signals were additive (**Fig. 2F**) and antigen-receptor signaling redirected STAT4 genomic binding (**Fig. 2G** and **Supp. Fig. 4C**). We speculated that differentially bound loci harbor distinct TF-binding motifs. Indeed, regions with an antigen-dependent decrease in STAT4 binding were relatively enriched for ETS, RUNX and STAT motifs, whereas regions with an antigen-dependent increase in STAT4 genomic binding were particularly enriched for AP-1 motifs (**Fig. 2H** and **Supp. Fig. 4D**). Together, antigen receptor signaling redirects STAT4 genomic binding away from ETS, RUNX and STAT motifs and towards AP-1 binding sites.

### AP-1 acts as a pioneer factor to enable AP-1/STAT4 cooperation

To further understand how regions containing AP-1 motifs increase STAT4 genomic binding, we performed STAT4 CUT&RUN with a pan-AP-1 inhibitor (AP1i) (**Supp. Fig. 5A and 5B, Fig. 3A-C**). Pre-incubation of NK cells with AP1i specifically reduced antigen-dependent STAT4 genomic binding in peaks containing AP-1 motifs (e.g. peak 2621 and 3254), but not in peaks without AP-1 motifs (**Fig. 3A** and **3B**, e.g peaks 2616 - 2619, 3252 - 3253). AP-1 factors have been proposed to act as pioneer factors regulating chromatin accessibility^26^. To test if antigen signaling alters chromatin accessibility via AP-1, we performed ATAC-seq of NK cells stimulated with antigen-receptor with and without AP1i (**Fig. 3D-E** and **Supp. Fig. 5C**). In addition, we conducted CUT&RUN for cJUN to directly visualize AP-1 binding (**Fig. 3D-E**, **Supp. Fig. 5D**), since cJUN was expressed higher than other AP-1 family members and induced by antigen stimulation in NK cells (**Supp. Fig. 5E**). Regions where AP-1 inhibition reduced STAT4 genomic binding showed AP1-dependent changes in chromatin accessibility and cJUN binding (**Fig. 3D-E, Supp. Fig. 5F**). Globally, regions that bound cJUN after antigen stimulation gained accessibility (**Fig. 3F** and **Supp. Fig. 5G**) and showed increased STAT4 genomic binding after subsequent IL-12 stimulation (**Fig. 3G** and **Supp. Fig. 5H**). Together, our findings suggest that antigen priming leads to AP-1 dependent chromatin changes that facilitate genomic relocation of STAT4 (**Fig. 3H**).

**Figure 3:**
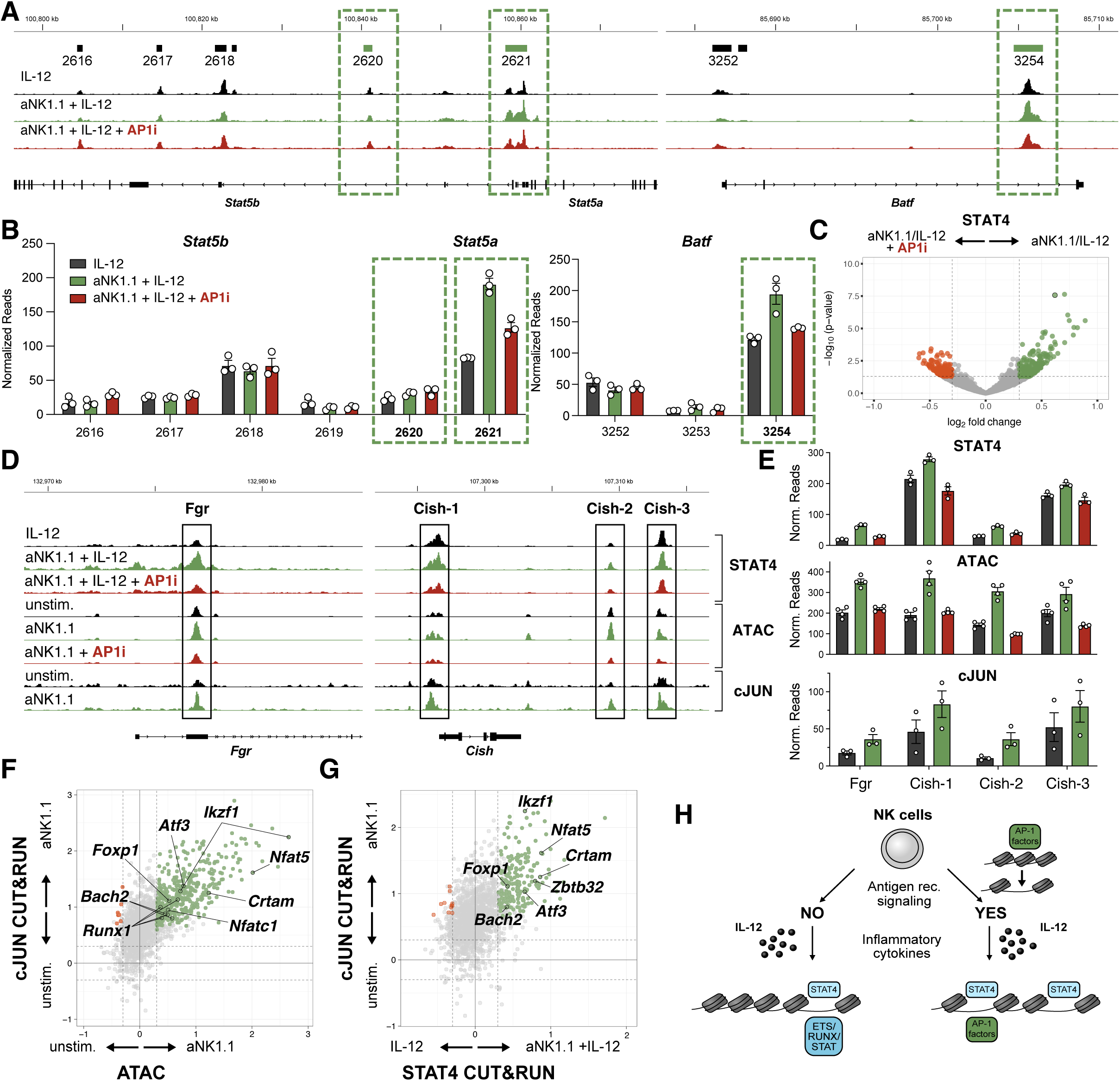
AP-1 acts as pioneer factor to redirect STAT4 genomic binding in adaptive NK cells. **A)** NK cells were stimulated with IL-12, aNK1.1 + IL-12, or AP1i + aNK1.1 + IL-12 (stimulation sequence: +/-AP1i ➔ +/- aNK1.1 ➔ +/- IL-12, also see **Supp. Fig. 5A** for experimental setup). Example normalized STAT4 CUT&RUN tracks averaged across all replicates are shown. Peaks are denoted as a bar above the tracks with a peak ID, and peaks with green bar and outlined with a green dotted line contain AP-1 motifs. **B)** Quantification of normalized read counts from peaks shown in A). Boxes with dotted line indicate peaks containing AP-1 motifs from A). **C)** Volcano plot of STAT4 genomic binding after aNK1.1 + IL-12 versus AP1i + aNK1.1 + IL-12 stimulation. Highlighted are regions with p <0.05 and log2foldchange > 0.3. **D)** Example peaks for STAT4 CUT&RUN, ATAC-seq and c-JUN CUT&RUN (normalized with flanking method) in the indicated conditions. **E)** Quantification of peaks shown in D). **F)** Scatterplot between the overlapping peaks of ATAC-seq and cJUN CUT&RUN for NK cells stimulated with aNK1.1 versus unstimulated control. **G)** Scatterplot between the overlapping peaks of STAT4 CUT&RUN and cJUN CUT&RUN for NK cells stimulated versus control in the respective conditions. **H)** Schematic of STAT4 genomic binding in antigen-primed and antigen-unprimed NK cells.

### Step-wise signal integration is required for an adaptive fate decision in NK cells

Our data indicates that antigen signaling redirects STAT4 genomic binding, suggesting that IL-12 signaling may differentially impact NK cells with or without the antigen receptor Ly49H. Indeed, whereas IL-12 signaling was crucial for efficient adaptive NK cell responses in the Ly49H^+^ subset (**Fig. 4A** and **Supp. Fig. 6A)**^17^, IL-12 signaling <u>impaired</u> expansion of Ly49H^-^ (= antigen receptor negative) NK cells (**Fig. 3B** and **Supp. Fig. 6B**). This inverse impact of IL-12 on antigen receptor expressing or negative cells was visible as early as day 4 PI, marking the beginning of adaptive NK cell responses (**Supp. Fig. 6C)**^27^. We hypothesized that in the absence of prior antigen signaling, IL-12 may lead to terminal instead of adaptive differentiation of NK cells, explaining the context-dependent impact on expansion. Indeed, without prior antigen receptor signaling, IL-12 impaired NK cell proliferation in *in vitro* assays (**Fig. 4C-F**), and pre-incubating NK cells with IL-12 completely abrogated their adaptive potential *in vivo* (**Fig. 4G**). Because IL-12 exposure in the absence of preceding antigen signals completely arrested proliferation in many NK cells, we attempted to find evidence for this proliferation arrest in our Nur77 system. Indeed, CTV labeling of co-transferred Nur77^high^ and Nur77^low^ (unprimed) NK cells revealed an accumulation of undivided NK cells within the transferred Nur77^low^ NK cell population (**Fig. 4H**).

**Figure 4:**
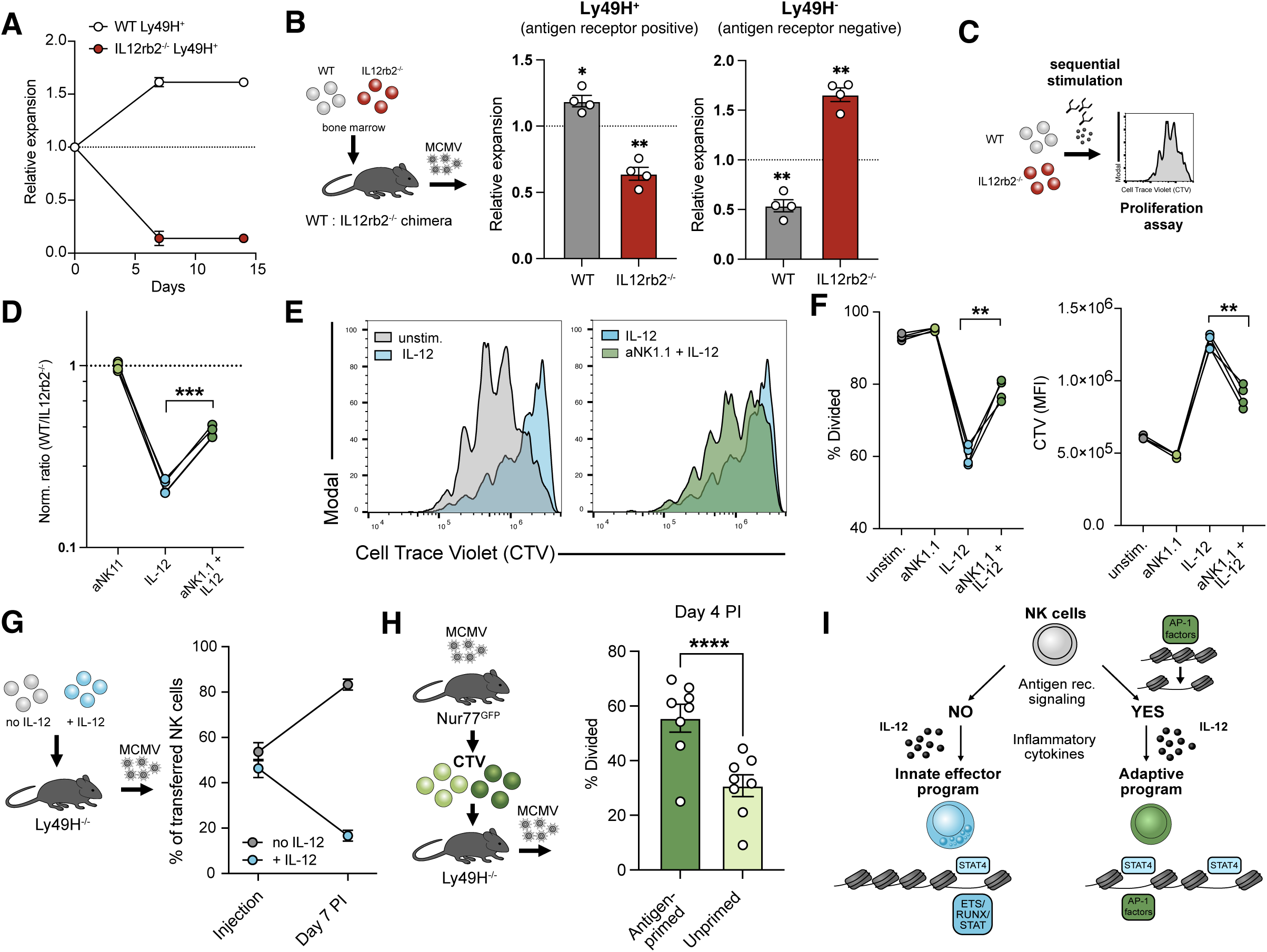
Preceding antigen signaling promotes innate versus adaptive fate decision in NK cells. **A)** Adoptive co-transfer of WT and *Il12rb2^-/-^* NK cells into Ly49H-deficient hosts and measurement of expansion following MCMV infection. **B)** Expansion of NK cells in competitive WT:*Il12rb2^-/-^* chimera on day 7 PI separated into Ly49H^+^ (antigen-receptor expressing) and Ly49H^-^ (antigen-receptor negative) NK cells. **C)** Schematic of Cell Trace Violet (CTV) dilution proliferation assay. **D)** Relative abundance of WT and *Il12rb2^-/-^* NK cells in IL-12, anti-NK1.1, or sequentially stimulated NK cells after 3 days of stimulation with IL-15. **E)** Representative histograms showing CTV dilution in IL-15 dependent proliferation assay. **F)** Quantification of divided cells and CTV MFI. **G)** Co-transfer of untreated or IL-12 pre-incubated NK cells into Ly49H^-/-^ mice infected with MCMV and measurement of expansion. **H)** Nur77^high^ (antigen-primed) and Nur77^low^ (unprimed) NK cells on day 1 PI from Nur77-GFP mice were isolated, stained with CTV and co-transferred into infection-matched recipients. Measurement of division on day 4 PI. **I)** Schematic of fate decision between innate effector program and adaptive program via differential integration of inflammatory signaling in antigen receptor positive and negative NK cells. Data in Fig. 4A, 4B), 4G) and 4H) are pooled from 2 individual experiments. Data in 4D) and 4F) are representative of 2 individual experiments. Error bars represent SEM. Significances are calculated as paired T-test (competitive chimera, co-transfers, or same biological replicates). **** p < 0.0001, *** p < 0.001, ** p < 0.01, * p < 0.05.

Together, our analysis of adaptive NK cell responses suggests that differential integration of inflammatory cytokine signals promotes distinct fate decisions in NK cells: without prior antigen signaling, STAT4 induces terminal differentiation of NK cells incapable of proliferation by binding to genomic regions shared with RUNX and ETS motifs (**Fig. 4I**). Sufficient antigen-receptor signaling redirects subsequent STAT4 genomic binding towards AP-1 binding sites and instead promotes adaptive programming of NK cells (**Fig. 4I**).

### Integration of inflammatory cytokine signals in CD8^+^ T cells depends on antigen strength

Previously, we have found shared epigenetic features in the adaptive programming of innate and adaptive lymphocytes^28^. Naïve CD8^+^ T cells do not express the IL-12 receptor until activation^29^, making it impossible to study the impact of IL-12 signaling in T cells that have not seen antigen. Thus, we instead hypothesized that in CD8^+^ T cells the strength of antigen signaling (rather than the presence or absence of antigen receptor) may be what impacts subsequent cytokine signaling. To test this idea, we generated retrogenic mice from WT or *Il12rb2^-/-^* stem cells transduced with a high- or low-affinity TCR from a previously generated library of SIINFEKL peptide-specific TCRs^30^ (**Supp. Fig. 7A**). When retrogenic T cells were transferred into mice infected with recombinant *Listeria monocytogenes* expressing SIINFEKL (*L.m.*-SIINFEKL), IL-12 promoted clonal expansion in high-avidity (B11) but not low-avidity (E8) CD8^+^ T cells (**Fig. 5A**). Comparing memory responses after challenge with the heterologous pathogen MVA-dE5R-OVA^31^, high-avidity T cells similarly showed a greater recall potential dependent on IL-12. In contrast, low-avidity T cells with intact IL-12 signaling had impaired memory responses (**Fig. 5B**).

**Figure 5:**
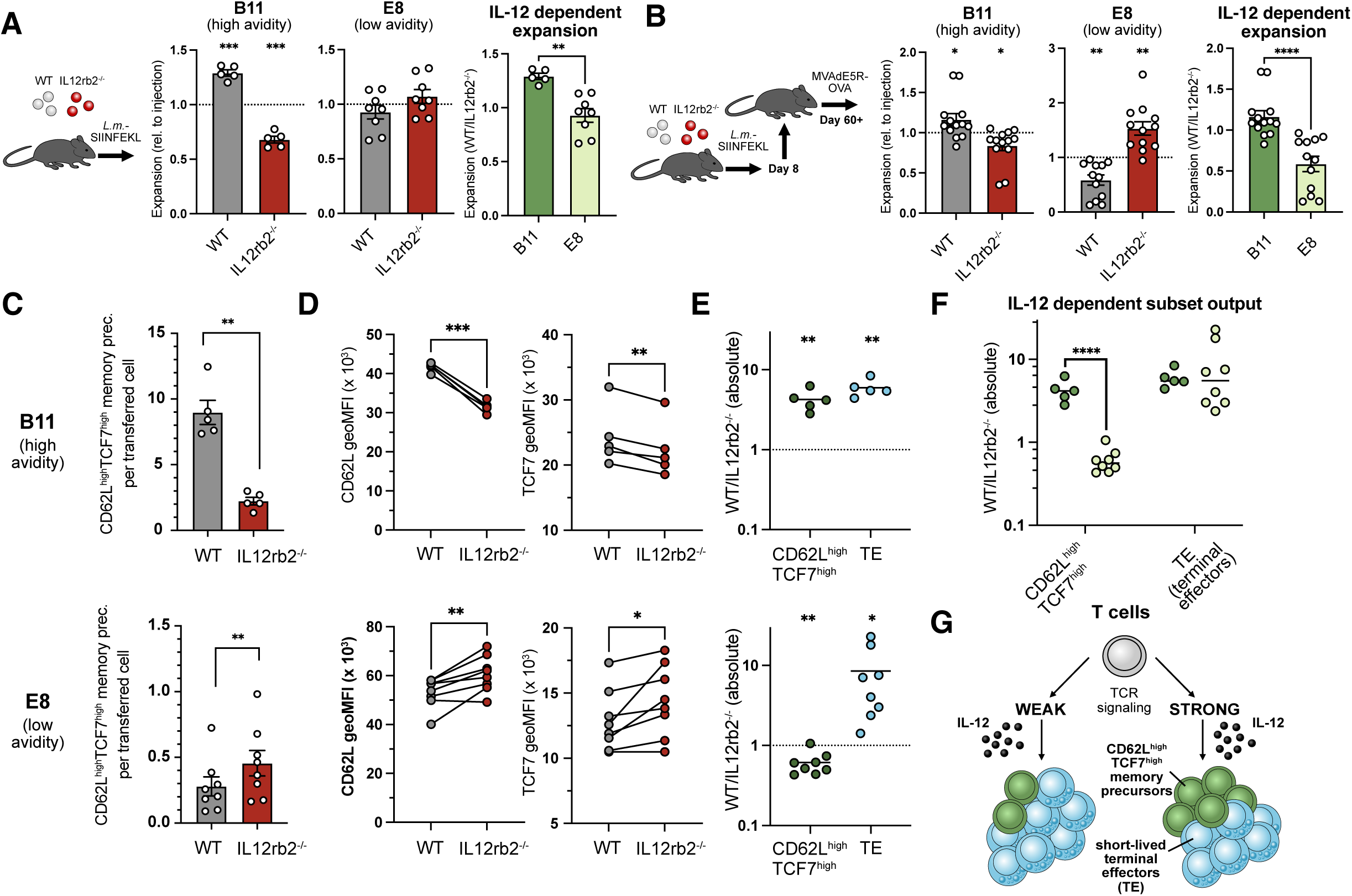
Avidity-dependent integration of inflammatory cytokine signaling drives CD8^+^ T cell differentiation. **A)** (Left) Acute expansion (day 8) of high-avidity (B11) and low-avidity (E8) WT and *Il12rb2^-/-^*CD8^+^ T cells in mice infected with *L.m*.-SIINFEKL. (Right) Comparison of WT/*IL12rb2^-/-^* ratios of absolute per-cell output for high- and low-avidity T cells during acute response. **B)** (Left) Recall expansion of high-avidity (B11) and low-avidity (E8) WT and *Il12rb2^-/-^*CD8^+^ T cells during recall response against MVAdE5R-OVA. Retransfer of whole splenocytes into secondary recipients on day 8 PI and recall infection after > 60 days post transfer. (Right) Comparison of WT/*Il12rb2^-/-^*ratios of absolute per-cell output for high- and low-avidity T cells. **C)** Comparison of WT and *Il12rb2^-/-^* CD62L^high^TCF7^high^ memory precursor output per transferred cell for high-avidity (top, B11) and low-avidity (bottom, E8) T cells during acute response against *L.m*.-SIINFEKL. **D)** CD62L and TCF7 MFIs of CD62L^+^ T cells derived from high-avidity (top, B11) and low-avidity (bottom, E8) WT and *Il12rb2^-/-^*CD8^+^ T cells. **E)** WT/*Il12rb2^-/-^* ratio of absolute cell output for CD62L^high^TCF7^high^ memory precursor and terminal effector (TE) cells for high-avidity (top, B11) and low-avidity (bottom, E8) T cells. **F)** Direct comparison of IL-12 dependent CD62L^high^TCF7^high^ memory precursor and terminal effector (TE) output for high-avidity (dark green, B11) and low-avidity (light green, E8) T cells. **G**) Schematic: antigen signal strength directs inflammation-dependent fate decision between memory and effector fates in high- and low-avidity CD8^+^ T cells. Data in 5A), 5C) – F) are representative of 2-3 independent similar experiments. Data in 5B) is pooled from 2 individual experiments. Error bars represent SEM. Significances are calculated using paired or unpaired T-test. **** p < 0.0001, *** p < 0.001, ** p < 0.01, * p < 0.05.

We hypothesized that the observed differences in memory potential were due to differential generation of memory precursor populations (**Supp. Fig. 7B**). Using different marker combinations, we confirmed previous studies that showed defective differentiation of terminal effector cells by *Il12rb2^-/-^* T cells^6,7^ (**Supp. Fig. 7C and 7D**). Comparing memory precursor populations, only high-avidity T cells showed an increase in differentiation into CD62L^high^ TCF7^high^ memory precursor cells^32,33^ (**Supp. Fig. 7D and 7E**). On average, WT high-avidity T cells generated far more CD62L^high^ TCF7^high^ memory precursor cells than *Il12rb2*^-/-^ counterparts (9.0 vs 2.0, **Fig. 5C, top**). However, among low-avidity T cells, WT cells generated less CD62L^high^ TCF7^high^ memory precursors than *Il12rb2*^-/-^ controls (0.28 vs. 0.45, **Fig. 4C, bottom**). In general, CD62L^+^ WT cells showed higher expression of CD62L and TCF7 than *Il12rb2*^-/-^ counterparts (**Fig. 5D, top**), whereas the opposite was observed in low-avidity T cells (**Fig. 5D, bottom**).

Quantifying the IL-12 dependent output of subsets (WT:*Il12rb2^-/-^*ratio), terminal effector cell output was similarly dependent on IL-12 signaling in both high- and low-avidity T cells (**Fig. 5E** and **5F**). CD62L^high^ TCF7^high^ memory precursor cells, however, were generated in an IL-12 dependent manner in high-but not low-avidity T cells. Thus, in CD8^+^ T cells, antigen receptor signal strength resulted in differential integration of subsequent inflammatory signals. This mechanism led to a preferential accumulation of CD62L^high^ TCF7^high^ memory precursors and increased memory potential in high-avidity T cells, whereas output of short-lived effector cells was not impacted by avidity (**Fig. 5G**).

### Step-wise signal integration is a conserved feature in innate and adaptive lymphocytes

Our data reveals a context- and antigen-dependent role for IL-12 in both NK cells and CD8^+^ T cells. To test whether antigen receptor signaling in T cells redirects STAT4 genomic binding as observed for NK cells, we set up an *in vitro* model of sequential stimulation. To mimic high-avidity and low-avidity TCR signaling, we used altered peptide ligands for SIINFEKL-specific OT-I T cells^34^, and titration of peptides revealed an avidity-dependent upregulation of the activation marker CD69 (**Fig. 6A**). Since CD8^+^ T cells do not express the IL-12 receptor until activation via the TCR^29^, we assessed the earliest timepoint at which CD8^+^ T cells can detect IL-12 signaling. A kinetic experiment revealed that both high- and low-avidity T cells first begin to show IL-12-dependent pSTAT4 at 12-16h following peptide stimulation (**Fig. 6B** and **6C**). Although memory T cells showed low levels of pSTAT4 with IL-12 stimulation alone, antigen re-stimulation substantially increased phosphorylation of STAT4, suggesting that both naïve and memory CD8^+^ T cells are similarly shielded from IL-12 signaling without prior antigen (re-)stimulation (**Supp. Fig. 8A**). Culturing high- and low-avidity stimulated T cells in presence or absence of IL-12, we found evidence of differential integration of inflammatory signals, recapitulating the phenotype observed *in vivo*: IL-12 signaling increased CD62L expression in high-avidity T cells (**Fig. 6D and Supp. Fig. 8B**). In contrast, CD62L expression level was diminished in low-avidity T cells after IL-12 stimulation.

**Figure 6:**
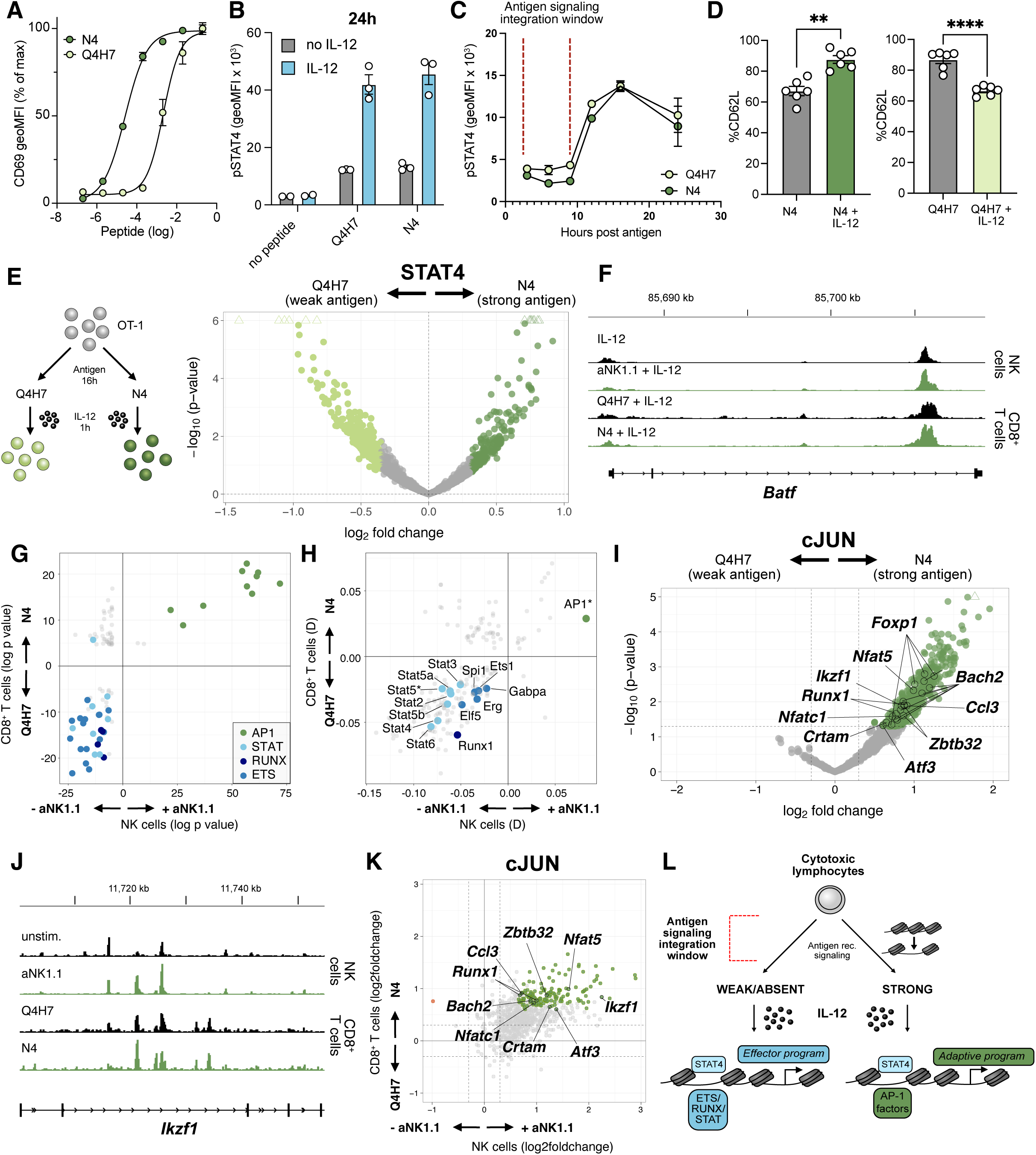
Sequential integration via STAT/AP-1 cooperation shape effector versus memory differentiation in cytotoxic lymphocytes. **A)** Peptide titration of SIINFEKL (N4, high-avidity) and SIIQFEQL (Q4H7, low-avidity)^34^ stimulated OT-1 T cells. **B)** pSTAT4 staining of IL-12 stimulated T cells after 24h of peptide stimulation. **C)** Time titration of pSTAT4 MFI after 1h of IL-12 stimulation of peptide stimulated OT-1 T cells. **C)** Flow cytometry analysis of CD62L expression levels after 3 days of high- and low-avidity peptide +/- IL-12. **E)** Volcano plot of STAT4 CUT&RUN differential binding analysis using median-of-ratios normalization. Light green indicates the top 20% peaks by absolute log2 fold change that increase in binding upon weak antigen compared to strong antigen stimulation. Dark green indicates the top 20% peaks by absolute log2 fold change that increases in binding upon strong antigen compared to weak antigen. Triangle shape indicates peaks above y axis limit. **F)** Tracks of STAT4 CUT&RUN at the *Batf* locus in NK cells and CD8^+^ T cells given indicated stimulations. **G)** Scatter plot of log P value in HOMER known motif analysis. The top 20% peaks denoted above, as well as their equivalent in NK cell analysis, were analyzed for enrichment for known motifs in HOMER database, with STAT4 binding atlas peaks as background. Results were filtered with HOMER’s default P value threshold (1e-1) and q value threshold of <= 0.05. Motifs comparatively enriched for weak/no antigen in the respective datasets were assigned negative values in the respective axes. Colors represent manual categorization of the motifs. **H)** Scatter plot of motif enrichment analysis D statistics from KS tests as described in Figure 2E between NK cells and CD8^+^ T cells. *AP-1 motif labelled here overlaps three motifs, Atf3, Jun and Fos::Jun. **I)** Volcano plot of cJUN CUT&RUN normalized with flanking method in CD8^+^ T cells stimulated with strong (N4) versus weak (Q4H7) antigen. **J)** Example tracks from c-JUN CUT&RUN in NK cells and CD8^+^ T cells in the indicated conditions. **K)** Scatterplot of cJUN differential binding dependent on antigen stimulation for CD8^+^ T cells and NK cells. **L)** Schematic: AP-1/STAT cooperation underlies antigen-dependent differential integration of inflammatory cytokine signaling in innate and adaptive lymphocytes to promote an adaptive/memory fate. Data in 6A) to 6D) are representative of two experiments. Error bars represent SEM. Significances were calculated using paired T-test (same biological replicates). **** p < 0.0001, *** p < 0.001, ** p < 0.01, * p < 0.05.

Confident that we can model avidity-dependent integration of inflammatory signals *in vitro*, we stimulated T cells with high- or low-avidity peptides followed by IL-12 and performed CUT&RUN to determine STAT4 binding (**Fig. 6E** and **6F**, **Supp. Fig. 8C**). To compare STAT4 genomic binding in CD8^+^ T cells and NK cells, we performed a comparative motif enrichment analysis. We found that, as observed for NK cells, strong antigen signaling redirected STAT4 away from binding sites containing STAT, RUNX and ETS towards AP-1 motifs in CD8^+^ T cells (**Fig. 6F-H**, **Supp. Fig. 8D**). Similar to NK cells, genomic binding of c-JUN was dependent on antigen signaling and was markedly enhanced in T cells stimulated with high-avidity peptide (**Fig. 6I**). Furthermore, CD8^+^ T cells and NK cells share genomic sites with increased c-JUN binding dependent on antigen (**Fig. 6J** and **6K**). This data suggests that in cytotoxic lymphocytes, a conserved and step-wise AP-1/STAT4 cooperation regulates their adaptive potential.

Altogether, our data suggests that sufficient antigen receptor signaling must occur <u>prior</u> to cytokine signaling to drive an optimal adaptive fate in cytotoxic lymphocytes. This stepwise integration is mediated through an antigen strength-dependent STAT/AP-1 cooperation which is conserved among lymphocytes, whereby STAT4 is redirected away from STAT, RUNX and ETS motifs (**Fig. 6L**).

## Discussion

The immune system counters pathogen invasion by fulfilling two critical tasks: 1) it must rapidly contain the pathogen and protect the host through a primary response and 2) it must select the best equipped cells for memory formation in preparation of a more robust secondary response against reinfection. In the midst of a primary response against a pathogen, the immune system cannot afford to be too selective, as even cells with low-avidity against a quickly replicating pathogen are preferable to cells that do not recognize the pathogen at all. However, for memory generation, amplifying the number of immune cells with high-avidity for the pathogen is desirable.

In our study, we find that the step-wise integration of antigen receptor and inflammatory cytokine signaling acts to optimize both tasks highlighted above. Whereas immune cells receiving no or inadequate antigen receptor signaling are driven into a terminal effector fate by inflammatory cytokines during the antiviral response, enhanced memory formation of high-avidity cells requires adequate antigen receptor signals prior to encountering inflammation. Previous studies have shown that low-avidity T cells are capable of effector generation^34^, and in fact generate effector cells earlier than high-avidity T cells^35^, which receive prolonged TCR signaling and stronger interactions with antigen-presenting DCs. Our study suggests an alternative or additional mechanism, by which the immune system may promote early effector differentiation of low-avidity T cells and NK cells: Inflammatory cytokines promote a terminal effector fate in “suboptimal” cells, while preferentially selecting high avidity immune cells for adaptive responses. Fittingly, DCs are the main source of IL-12 during infection^36,37^, and could thereby directly govern this fate decision by providing both antigen and cytokine signals.

Previous studies have established that IFN signaling can suppress the proliferative capacity of naïve CD8^+^ T cells^38,39^, while IFN shortly before or after antigen signaling can promote adaptive responses^40^. The proposed mechanism for this effect is a direct inhibitory effect of IFN on cell proliferation, although this mechanism has not been fully established. Interestingly, another proposed mechanism is that antigen receptor signaling suppresses STAT1 while promoting STAT4 activation^41^. In contrast to these models, here we propose that STAT4 activation via IL-12 impairs adaptive responses of NK cells via an epigenetic mechanism; however, if the NK cell first receives an antigen signal, STAT4 is redirected to instead promote an adaptive program. Future studies will shed light on how STAT1 versus STAT4 shape immune cell differentiation. Another interesting question is how antigen and IL-12 signaling shape the integration of subsequent signals. Prior studies in T cells have suggested that regulation of IL-2 signaling via IL-12 could contribute to the pro-proliferative effect of IL-12 in T cells^42,43^. Considering the epigenetic regulation of Il2ra and STAT5 in our data, this represents a likely downstream mechanism of antigen-dependent STAT4 redirection, the mechanism proposed in our study.

Although antigen-receptor signaling represents a main activator of AP-1 in lymphocytes, it is possible that additional stimuli can sufficiently activate AP-1 to redirect STAT4 genomic binding and facilitate adaptive programming. An obvious candidate is IL-18, which is often used together with IL-12 and IL-15 to generate adaptive-like NK cells *in vitro*^44^. Indeed, IL-18 has been shown to activate downstream AP-1^45,46^. Whether IL-18 drives similar or distinct AP-1 activation compared to activating receptors remains to be determined.

Interactions of STAT and AP-1 transcription factors have been proposed and observed in various immunological and non-immunological cell types. However, the main STAT transcription factor found to co-operate with AP-1 is STAT3: an AP-1/STAT3 cooperation has been found to regulate inflammatory memory of epithelial cells^47^, and influence the malignant potential of cancer^48^. In CD8^+^ T cells, STAT3 has been found to bind to genomic regions harboring AP-1, RUNX and ETS motifs^49^, the same motifs we identify here to be bound by STAT4. AP-1, ETS and RUNX transcription factors have been shown to affect CD8^+^ T cell^50–52^ and NK cell differentiation^53–55^. A recent study proposed STAT4-AP1 cooperation as a requirement for mediating the canonical genomic binding of STAT4 in general^56^. We observe that antigen signaling directs STAT4 away from RUNX and ETS sites, and instead promotes a STAT/AP-1 cooperation. Our findings suggest an additional layer of complexity whereby antigen signaling affects the degree to which inflammatory cytokine signaling interacts with distinct transcription factor networks to shape immune cell differentiation.

Beyond infection, our proposed signal integration model could prove crucial in other diseases such as autoimmunity and cancer^57,58^. In autoimmunity, it has been an outstanding question why inflammatory cytokines can have both protective and detrimental roles^59^. Our study offers a possible explanation. Autoimmunity can be elicited by T cells with low-avidity towards self-peptides that escape negative selection and become activated during inflammation or infection^60^. We suggest that STAT4 may act as a negative regulator for adaptive programming of low-avidity T cells; and once autoimmunity has been established, inflammatory cytokines could then promote expansion of these autoreactive cells.

In cancer, priming of tumor-reactive T cells in lymph nodes is characterized by decreased cytokine signaling compared to priming during infection^61^. At the same time, different tumors vary substantially in the amount of local immune activation, and immunologically “hot” tumors respond better to treatments such as immune checkpoint blockade^62^. Our study suggests that a lack of pro-inflammatory cytokines during priming in tumor-draining lymph nodes may have distinct effects on high- versus low-avidity T cells. Whereas high-avidity T cells will remain below their potential for mounting adaptive responses in the absence of inflammation, low-avidity T cells may in contrast be protected from inflammation-induced terminal differentiation. Because high-avidity T cells are often exhausted in tumor settings^63^, the targeting of low-avidity T cells may represent a promising strategy for cancer immunotherapy.

In conclusion, our work highlights a shared molecular mechanism underlying innate and adaptive lymphocyte differentiation, prompting new questions about optimal lymphocyte engineering, the context-dependent role of cytokines, and the relationship between the innate and adaptive immune systems.

## Acknowledgments

We thank past and present members of the Sun lab for helpful discussions and support. We thank Dr. Andri Lemarquis, Dr. Sasha Rudensky, Dr. Andrea Schietinger and Dr. Kilian Schober for input and/or critically reading the manuscript. Sequencing was performed by the Integrated Genomics Operation Core which is funded by the NCI Cancer Center Support Grant (CCSG, P30 CA08748), Cycle for Survival, and the Marie-Josée and Henry R. Kravis Center for Molecular Oncology. S.G. is the recipient of a CRI / Donald J. Gogel Postdoctoral Fellowship (CRI Award CR13934), was supported by the Deutsche Forschungsgemeinschaft (DFG, Award number GR5503/1-1) and NIAID (NIH) under award number K99AI180360. A.M.M. was supported by the CRI as a CRI/Amgen Fellow. M.O. was supported by the NIH T32 Predoctoral Training Grant (T32 AI134632-05) and the NIH F31 Ruth L. Kirschstein Predoctoral Fellowship from the National Institute of Allergy and Infectious Diseases (F31AI178958). S.X.F. was supported by a Medical Scientist Training Program grant from the National Institute of General Medical Sciences of the National Institutes of Health under award number T32GM152349 to the Weill Cornell/Rockefeller/Sloan Kettering Tri-Institutional MD-PhD Program. J.C.S. was supported by the Ludwig Center for Cancer Immunotherapy, the American Cancer Society, the Burroughs Wellcome Fund, and the NIH (AI100874, AI130043, AI155558, and P30CA008748).

## Methods

### Mice

All mice used in this study were housed and bred under specific-pathogen-free conditions with food and water in 12-h light–dark cycles at 72 °F with 30–70% humidity at Memorial Sloan Kettering Cancer Center and handled in accordance with the guidelines of the Institutional Animal Care and Use Committee. The following mouse strains were used in this study: C57BL/6 (CD45.2), C57BL/6 CD45.1 (CD45.1), C57BL/6 CD45.1×CD45.2, *Il12rb2^−/−^*, Klra8^−/−^ CD45.1xCD45.2 (Ly49H-deficient) and Ncr1-GFP and Nr4a1-GFP. Experiments were conducted using 8–10-week-old mice or 8–16 weeks post-transplant mixed bone-marrow chimeric mice and all experiments were conducted using age- and sex-matched mice in accordance with approved institutional protocols.

### MCMV virus preparation

MCMV (Smith strain) was serially passaged through BALB/c hosts three times and then salivary gland viral stocks were prepared with a homogenizer for dissociating the salivary glands of infected mice 3 weeks after infection.

### Mixed bone-marrow chimeras

Mixed bone-marrow chimeric (mBMC) mice were generated by lethally irradiating (950 cGy) C57BL/6 CD45.1×CD45.2 animals and reconstituting with a 1:1 mixture of bone-marrow cells from WT (CD45.1) and *Il12rb2^−/−^* (CD45.2) mice. Hosts were co-injected with anti-NK1.1 (PK136) to deplete any residual donor and host NK cells. Residual CD45.1^+^CD45.2^+^ host NK cells were excluded from all analyses. To reduce the effects of maturation, Ly49H^-^ NK cells were gated for positivity of Ly49D, an activating receptor not recognizing MCMV but instead by allogenic MHC molecules^65^.

### Infections of Nr4a1-GFP mice and mixed bone marrow chimera

Chimerism in WT: Il12rb2-/- chimera was assessed by blood staining 1-2 weeks prior to the experiment. Nr4a1-GFP mice and bone marrow chimera were infected i.p. with 5 x 10^3^ PFU of MCMV and analyzed at the respective time points using flow cytometry.

### Isolation of mouse NK cells and flow cytometry

Spleens were dissociated with glass slides and filtered through a 100-μm cell strainer. Flow cytometry experiments were analyzed using a Cytek Aurora (Cytek Biosciences). Cell sorting was performed using a BD Aria II cytometers (BD Biosciences). Before cell sorting, NK cells were enriched by incubating whole splenocytes with the following antibodies at 10 μg/ml: CD3ε (Clone 17A2), CD4 (Clone GK1.5), CD8 (Clone 2.43), Ter119 (Clone TER-119), CD19 (Clone 1D3), Ly6G (Clone 1A8) (BioXCell). After washing, cells were incubated with goat anti-rat beads (QIAGEN, cat. no. 310107). For intracellular staining, cells were fixed and permeablizied using eBioscience Intracellular Fix & Perm Buffer Set (Thermo Fisher, cat. no. 88-8824-00). For *ex vivo* RNAseq analysis in Fig. 1G, Ly49H^-^ NK cells were gated for positivity of Ly49D to reduce the effects of maturation.

### Cell culture for *in vitro* experiments

NK cells and T cells were cultured in complete IMDM (10% FBS, 1× L-glutamine, 1× sodium pyruvate, 1× BME, 1× MEM-NAA and 25 mM HEPES). For cytokine stimulations, we used 50 ng/ml mouse IL-15 (Peprotech, cat. no. 210-15), 100U/ml IFNa (R&D, cat. 12105) and 20 ng/ml mouse IL-12 (R&D Systems, cat. no. 419-ML-050) depending on the experiment.

### In vitro receptor and cytokine stimulation for sequencing of NK cells

High-binding 96-well flat-bottom plates were coated with 100 μl per well of 20 μg/ml anti-NK1.1 (Clone PK136, BioLegend, cat. 108759) in PBS at 4 °C overnight. To prevent antigen-receptor signaling in negative controls, we used negative enrichment via magnetic beads and Ncr1-GFP mice. Splenic NK cells from Ncr1-GFP mice were purified using the NK Cell Isolation Kit (Miltenyi, cat. 130-115-818). Depending on the condition, NK cells were stimulated for 1-3h in 96 wells coated with anti-NK1.1 or in uncoated wells. After 1-3h, NK cells were either sorted for GFP positivity and directly used for sequencing or stimulated for 3h with IL-12 (10ng/ml). RNA-seq was performed after 3h + 3h of stimulation. ATAC-seq was performed after 3h or 6h (Supp. Fig. 3C). STAT1, STAT4 and STAT5 CUT&RUN was performed after 3h of aNK1.1 stimulation and 1h of IFNa, IL-12 or IL-15 stimulation. For controls, cells were incubated for 3h in uncoated wells with full media and 1h of stimulation (cytokines only). cJUN CUT&RUN was performed after 3h of antigen stimulation. For AP1 inhibition, cells were incubated for 1h with 50uM pan-AP1 inhibitor (SR 11302, Tocris, cat. 2476) or DMSO as control, washed off and stimulated according to the respective experiment.

### CTV dilution and proliferation assay

Splenocytes from CD45.1/.1 (WT1), CD45.1/.2 (WT2) and Il12rb2-/- CD45.2/.2 (Il12rb2-/-) were pooled and enriched using the NK Cell Isolation Kit (Miltenyi, Cat. 130-115-818). For CTV assays, cells were stained with CellTrace Violet (CTV) according to the manufacturer’s instructions (Thermo Scientific, cat. C34557). NK cells were then stimulated in aNK1.1. coated plates and stimulated with cytokines according to the respective experiment, or directly pooled and injected (Fig. 4H). Sequential stimulation was performed as 3h of aNK1.1 and 3h of IL-12 stimulation before adding of IL-15 for overnight incubation. For CTV assays, NK cells were kept after sequential stimulation in aNK1.1 coated 96-well U-bottom plates overnight and then re-plated in 96-well U-Bottom plates without aNK1.1 coating for an additional 2 days.

### Generation of retrogenic mice

Retrogenic mice were generated as described in^30^. Briefly, HSCs were sorted by staining with Sca-1 antibodies and expanded in DMEM containing 10%FCS, 1% Penicillin/Streptomycin, 1% L-Gln, 2ng/ml mIL-3 (Peprotech), 50ng/ml mIL-6 (Peprotech) and 50ng/ml mSCF (Peprotech) for 3 days. Retronectin coated plates were coated with retrovirus generated with HEK cells transfected with mP71 vectors and pl-ECO packaging vectors (using Lipofectamine™ LTX Reagent with PLUS™ Reagent) by centrifugation at 3000g for 2h. Cells were spinoculated for 90min at 800g, cultured for 2 additional days and injected into mice irradiated with 950 cGy prior to injection of HSCs.

### CD8^+^ T cell peptide stimulation

Whole splenocytes from OT-1 mice (C57BL/6-Tg(TcraTcrb)1100Mjb/J) were seeded at a concentration of 400.000/well of a 96 well plate. Peptide (SIINFEKL = N4 or SIIQFEHL = Q4H7) was serially diluted and added. IL-12 was added at a final concentration of 10ng/ml where applicable. For CUT&RUNs and CD62L staining analysis, 10ng/ml of peptide were used.

### T cell transfer and listeria infections

Frequency of retrogenic T cells was assessed by multimer staining (SIINFEKL multimer, NIH Core Facility). T cells were sorted as CD8^+^ CD44^low^ (naïve cells, acute expansion) or CD8^+^ CD44^high^ (activated cells, day 8 retransfer for recall experiment) and injected into recipients i.v. Listeria-OVA was cultured in 5ml of BHI media (BD Biosciences) with Erythromycin overnight. 50ul of overnight culture was inoculated into 5ml of fresh BHI + Erythromycin and grown to an optical density (OD600) of 0.9 – 1.1. OD of 1.0 was set to correspond to a bacteria concentration of 2 x 10^8^ CFU/ml. 5.000 CFU (transfer experiments) or 10.000 CFU (direct infection of retrogenics) were injected per mouse in 200ul of PBS i.v one day after T cell injection. For acute analysis, whole spleen was pre-enriched afte red blood cell lysis using negative depletion of B cells and Granulocytes (anti-CD19, anti-Ly6g). Counting beads were used to calculate absolute cell number per spleen. In recall experiments, blood was collected and analyzed via flow cytometry after red blood cell lysis. Mice where one of the transferred populations had less than 3 total events were excluded.

### MVA-**Δ**E5R-OVA infections

It was shown previously that deletion of E5R gene, which encodes an inhibitor of the DNA sensor, cyclic GMP-AMP synthase (cGAS), from the MVA genome, enhances antigen-specific CD8^+^ T cell responses^31^. MVAΔE5R-OVA virus expressing mCherry were generated as described^31^. The virus was propagated in BHK-21 cells and purified through a 36% sucrose cushion. Viral titers were determined using BHK-21 cells. Mice were infected with 1 x 10^7^ PFU i.p.

### Mouse single-cell RNA-sequencing and analysis

Splenic Ly49H^+^ NK cells (CD3/TCRb/CD19−NK1.1^+^CD49b^+^Ly49H^+^) from uninfected or MCMV-infected WT C57BL/6 mice were sorted as described and stained with barcoded antibodies (Total-Seq B, BioLegend). After hash-staining, NK cells were pooled and the single-cell RNA-seq from these pooled FACS-sorted cell suspensions was performed on a Chromium instrument (10x Genomics) following the user guide manual for 3′ v.3.1. In brief, FACS-sorted cells were washed once with PBS containing 1% bovine serum albumin (BSA) and resuspended in 1× PBS containing 1% BSA to a final concentration of 700–1,300 cells per μl. The viability of cells was above 80%, as confirmed with 0.2% (w/v) Trypan blue staining (Countess II). Cells were captured in droplets. Following reverse transcription and cell barcoding in droplets, emulsions were broken and cDNA-purified using Dynabeads MyOne SILANE (Thermo Fisher, cat. 37002D) followed by PCR amplification as per the manual instructions. Samples were multiplexed together on one lane of 10x Chromium (using Hash Tag Oligonucleotides (HTOs)). Final libraries were sequenced on Illumina NovaSeq6000 S4 platform (R1, 28 cycles; i7, 8 cycles; and R2, 90 cycles). The cell–gene count matrix was constructed using the 10x Cell Ranger (v7.1.0) pipeline based on mm10 2020-A reference. Viable cells were identified on the basis of library size and complexity, whereas cells with >20% of transcripts derived from mitochondria were excluded from further analysis. After mitochondrial and doublet cleanup, the raw count matrix was normalized by median library size normalization followed by log transformation. The Louvain algorithm with k = 30 was used to perform clustering and Euclidean distance was used as the metric to construct a nearest-neighbor. ‘Receptor signature’ was constructed from bulk RNA-seq data of naive NK cells stimulated with anti-NK1.1 for 3 hours *in vitro*, while ‘Cytokine signature’ was obtained from GSEA mouse REACTOME gene set (R-MMU-1280215). All downstream analysis was done using scanpy (v1.9.0).

### Bulk RNA sequencing

For data generated in this study, RNA was isolated from sorted cells and/or stimulated cells as described using PicoPure RNA Isolation Kit (ThermoFisher Scientific, cat. KIT0214). For *ex vivo* RNAseq analysis in Fig. 1F, Ly49H^-^ NK cells were gated for positivity of Ly49D to reduce the effects of maturation. After RiboGreen quantification and quality control by Agilent BioAnalyzer. RNA was then amplified using SMART-Seq v.4 Ultra Low Input RNA kit (Clontech, cat. 63488). Subsequently, amplified complementary DNA was used to prepare libraries with KAPA HyperPrep Kit (Kapa Biosystems, cat. KK8504). Samples were barcoded and ran on NovaSeq6000 in a 100bp/100bp paired-end run.

### Bulk ATAC sequencing

ATAC-seq was performed as described previously^66^. Briefly, sorted, stimulated cells were washed in cold PBS and lysed. The transposition reaction was incubated at 37 °C for 30 min. DNA was cleaned with the MinElute PCR Purification kit (QIAGEN, cat. 28004) and material was amplified for 5 cycles. After evaluation by real-time PCR, 8–11 additional PCR cycles were performed. The final product was cleaned by AMPure XP beads (Beckman Coulter catalog no. A63882) at a 1× ratio, and size selection was performed at a 0.5× ratio. Libraries were sequenced on a NovaSeq6000 in a 100 bp/100 bp paired-end run, using the NovaSeq6000 S2 or S4 Reagent kit (Illumina).

### Transcription factor CUT&RUN

For transcription factor CUT&RUN, 200,000 - 500,000 sorted total NK cells and CD8^+^ T cells were used. T cells were sorted for homogeneous expression of activation markers (CD69 and additionally CD44 for CD8^+^ T cells). Cells were light fixated with 0.1% PFA and fixated in antibody buffer (1X eBioscience Perm/Wash Buffer, 1X Roche cOmplete EDTA-free Protease Inhibitor, 0.5 uM Spermidine, + 2uM EDTA in H_2_O) prior to “staining” with polyclonal anti-STAT4 (ThermoFisher, cat. 71-4500, dilution 1/100), polyclonal anti-STAT1 (Proteintech, 10144-2-AP, dilution 1/100), polyclonal anti-STAT5b (ThermoFisher, cat 71-2500, 1/100) or cJUN (Cell Signaling Cat. 9165T, dilution 1/100) antibodies. Upon antibody incubation, cells were washed twice with Buffer 1 (1X eBioscience Perm/Wash Buffer, 1X Roche cOmplete EDTA-free Protease Inhibitor, 0.5 uM Spermidine in H_2_O) and resuspended in 50ul of Buffer 1 + 1X pA/G-MNase (Cell Signaling, cat. 57813) and incubated for 1 hour at 4°C. Cells were washed with Buffer 2 (0.05% w/v Saponin, 1X Roche cOmplete EDTA-free Protease Inhibitor, 0.5 uM Spermidine in 1X PBS) three times. After washing, Calcium Buffer (Buffer 2 + 2uM of CaCl_2_) was used to resuspend the cells for 30 mins at 4°C to activate the pA/G-MNase reaction, and equal volume of 2X STOP Buffer (Buffer 2 + 20uM EDTA + 4uM EGTA) was added along with 1 pg of *Saccharomyces cerevisiae* spike-in DNA (Cell Signaling, cat. 29987). Samples were incubated for 15 mins at 37°C, spun down at 18500g for 5min and supernatant was collected. DNA fragements were digested with Proteinase K (Thermo Fisher, cat. EO0491) under addition of 0.1% SDS for 4h at 65°C. DNA was isolated and purified using Qiagen MinElute Kit according to manufacturer’s protocol and subjected to library amplification^67,68^.

### RNA-seq, ChIP-seq, CUT&RUN, and ATAC-seq data processing

Data processing methods for published RNA-seq, ChIP-seq, and ATAC-seq datasets have been previously described^21,54^. For RNA-seq dataset generated in this study, transcript quantification was based on the mm10 University of California, Santa Cruz (UCSC) Known Gene models and performed using the quasi-mapping-based mode of Salmon (v0.13.1) correcting for potential GC bias. Transcript was summarized to gene level using tximport (v1.10.1). For ATACseq and CUT&RUN datasets found in this study, paired reads were trimmed for adaptors and removal of low-quality reads by using Trimmomatic (version 0.39) and aligned to the mm10 reference genome using Bowtie 2 (v2.4.1). ATACseq dataset was subjected to Tn5 correction prior to alignment. Upon alignment, peaks were called using MACS2 (v2.2.7.1) with input samples as a control using narrow peak parameters with cutoff-analysis -p 1e-5 --keep-dup all -B -- SPMR. For ATAC-seq data, reproducible peaks showing an IDR value of 0.05 or less in each condition were retained, aggregated across the experiment and merged via union generate the final atlas, and annotated with the UCSC Known Gene model. Reads were mapped to the final atlas and counted with the summarizeOverlaps function from the GenomicAlignment package (v1.34.1). For CUT&RUN data, peak atlas for each antibody targets was generated by merging via union of peaks called by each sample after filtering out bottom 25% of peaks by MACS2 calculated qValue. Resulting peak atlas was further filtered to only retain peaks that were called by two or more replicates within each sample group, as well as peaks that do not overlap the Encode’s DAC Exclusion List Regions (ENCFF547MET). CUT&RUN data was normalized with two methods as indicated in figure legends. Median-of-ratios method utilized the default DESeq2 (v1.46.0) parameters to generate size factor values for normalization, with the counts matrix of raw counts of the atlas peaks as input. For 3kb atlas flanking region method, 3kb of 5’ and 3’ flanking regions of the peak atlas were generated, only keeping the flanking regions that do not overlap with other atlas peaks. ‘estimateSizeFactors’ function from the DESeq2 was then used on the counts from these filtered flanking regions to generate size factor values for normalization. The rationale for using the median-of-ratios method for the STAT targets is to visualize genomic redistribution, whereas c-JUN showed a strong dependence on antigen stimulation and was therefore analyzed using the flanking method.

### Motif analysis

For motif enrichment analysis, each peak in the atlas was assessed using Find Individual Motif Occurrences (FIMO) from the MEME Suite (v5.4.1) against all motifs (224) from JASPAR CORE(v2024) mus musculus database. Peaks with antigen-dependent increase and decrease in STAT4 binding were compared with the total STAT4 atlas to examine motif enrichment using one-sided Kolmogorov-Smirnov (KS) test. The KS test statistic D is shown on the y axis and the proportion of regions associated with the motif is shown on the x axis. The odds ratio (frequency of the motif in increase or decrease group divided by its frequency in the entire atlas) was used to assign the motif to either left (regions with antigen-dependent decrease, negative value), or right (regions with antigen-dependent decrease, positive value) in x axis. Enrichment for known motifs was assessed using HOMER (v4.11) findMotifsGenome.pl with arguments mm10 -size given -len 6,8,10,12 -mset vertebrates -mask, with background set as other peaks in the respective STAT4 peak atlas.

### Downstream analyses of bulk RNA-seq and ATAC-seq

For data generated in this study, differential analyses were executed with DESeq2 (v1.38.3) using UCSC Known Gene models as reference annotations. For RNA-seq or ATAC-seq, genes or peaks are considered differential if they showed p-adjusted value of < 0.05, adjusted for multiple hypothesis correction. Motif enrichment analysis was conducted using HOMER algorithm (HOMER v.4.10) using regions that showed differential accessibility (p_adj_ < 0.05 & |log2FoldChange| > 0.5) found between conditions with parameter ‘–size given –len 6,8,10,12,15 –mset vertebrates –mast’ for de novo motif analysis. For correlation analyses, spearman coefficients were calculated using log_2_FC modeled by DESeq2 when comparing between conditions on either shared differential features (genes or peaks) for each condition, or all shared features.

### Visualization of RNA-seq, CUT&RUN and ATAC-seq

For peak-centered heatmaps and tracks, BAM files were converted to bigwig files using bamCoverage function and scaled based on sizeFactor modeled by DESeq2 or stated methods. Scaled bigwig files from each replicate per condition were then averaged using bigwigAverage function from deepTools (v3.5.4). All heatmaps from CUT&RUN and ATAC-seq experiments were plotted using EnrichedHeatmap (v1.28.1), while heatmaps from RNA-seq data were plotted using ComplexHeatmap (v2.14.0). All genomic tracks were either visualized by GViz (v1.42.1) or IGV (v2.9.4).

---

**Supp. Fig. 1:**
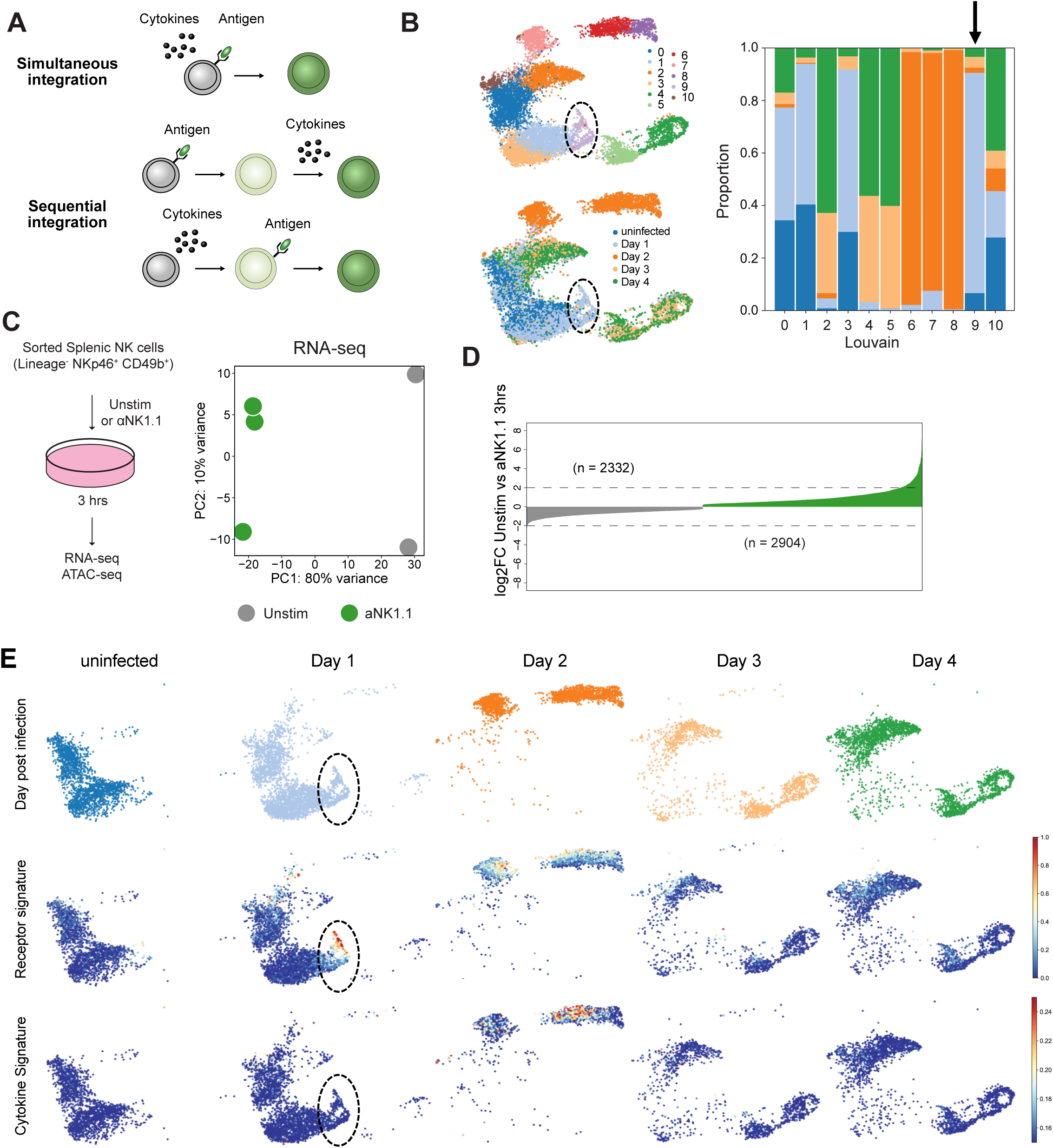
**A)** Concept of combinatorial signal integration and timed signal integration. **B)** (Left) Louvain clusters and hashtags identifying single-cell transcriptomes from different days PI. (Right) Proportion of single-cell transcriptomes from different days in Louvain clusters. Cluster 9 highlighted with arrow (enriched for day 1 but not uninfected) **C)** Schematic and PCA of RNAseq of antigen-receptor stimulated NK cells. **D)** Waterfall plot of log2fold change in gene expression after antigen receptor signaling. Top genes were used to assess antigen-receptor score. **E)** Full dataset of scRNA-seq of MCMV-reactive NK cells including days 3 and 4. Activating receptor signature and cytokine signature.

**Supp. Fig. 2:**
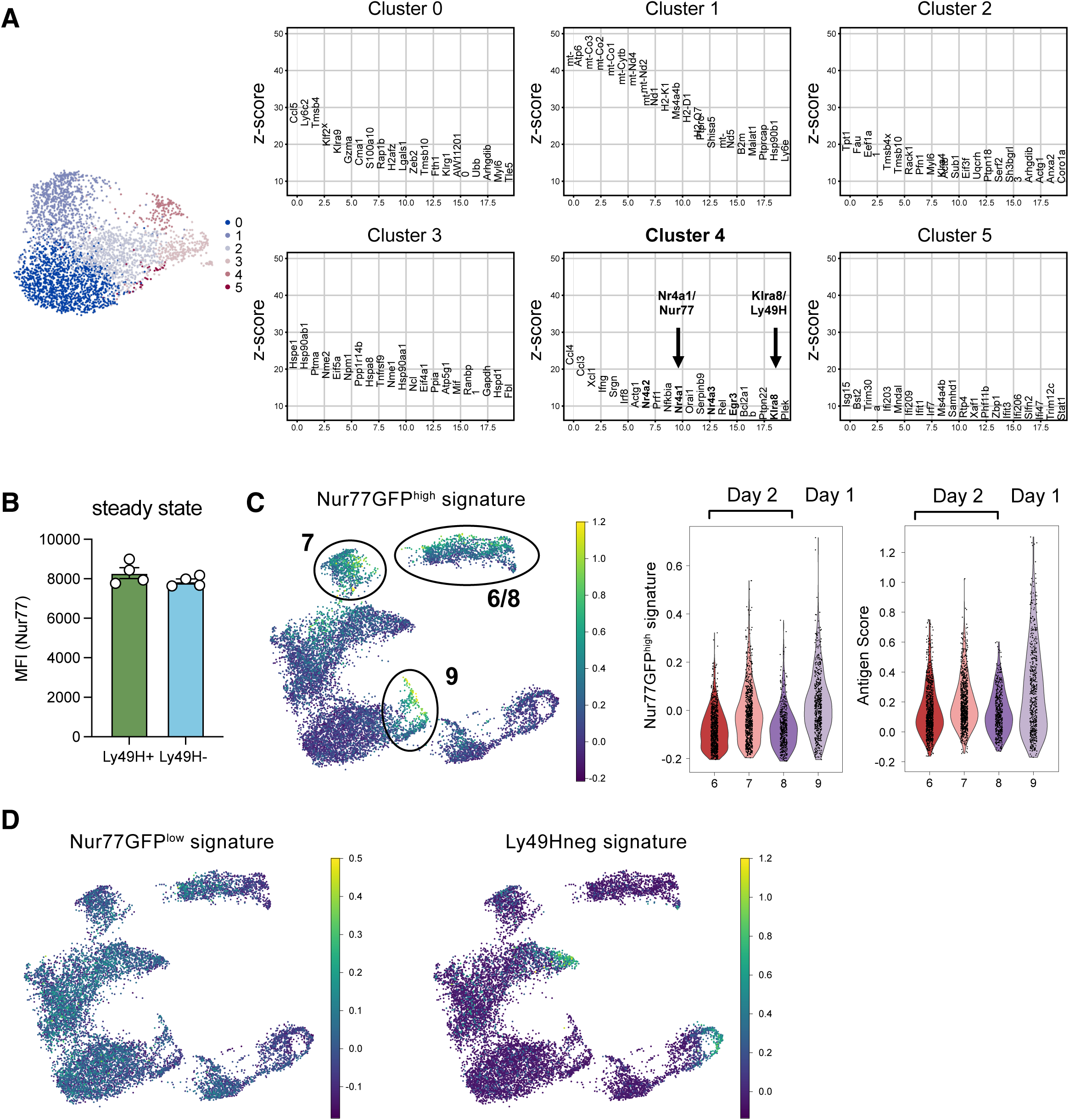
**A)** Ranking of genes in Louvain clusters on day 1 PI. Nr4a1/Nur77 and Klra8/Ly49H highlighted with arrows. **B)** Nur77^GFP^ MFI in Ly49H^+^ and Ly49H^-^ NK cells under steady state conditions. **C)** Gene signatures from pairwise comparisons against a common background were generated for the Nur77GFP RNAseq dataset in Fig. 1F-G. (left) The Nur77GFP^high^ signature was plotted in UMAP projection of scRNAseq from Fig. 1A-B. Highlighted are the approximate positions of clusters, for exact annotation refer to Fig. 1A and Supp. Fig. 1B. (right) violin plots for Nur77GFP^high^ signature and Antigen Score for clusters 6-9. Cluster 7 may represent antigen primed NK cells on day 2 PI. **D)** Nur77GFP^low^ and Ly49H^neg^ signature from data in Fig. 1F-G was plotted in UMAP projection of scRNAseq from Fig. 1A-B.

**Supp. Fig. 3:**
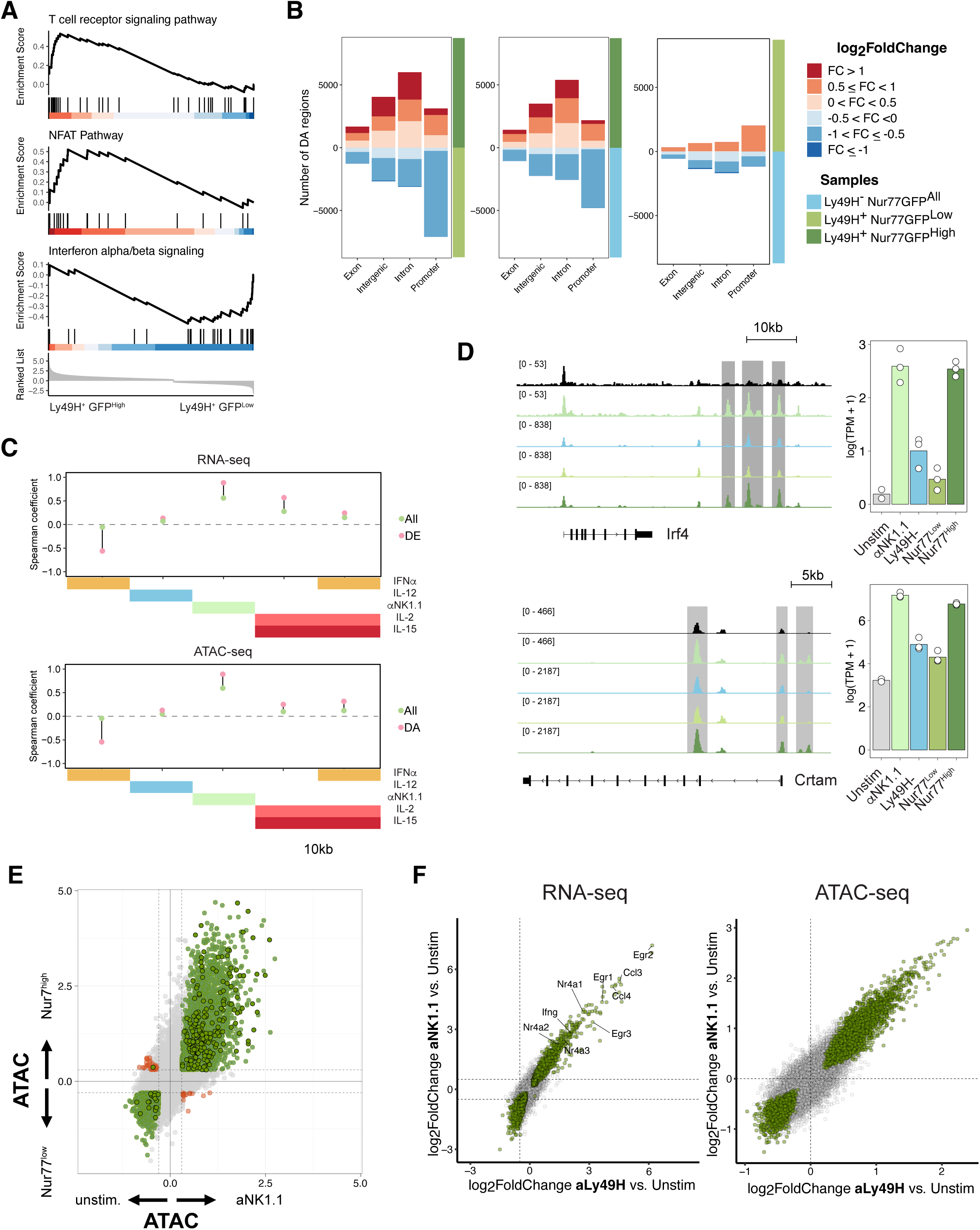
**A)** Pathway analysis of differentially expressed genes between Ly49H^+^ Nur77-GFP^high^ and Ly49H^+^ Nur77-GFP^low^ NK cells. **B)** Quantification and peak type analysis of differentially accessible regions in ATAC-seq of Ly49H^+^ Nur77-GFP^high^, Ly49H^+^ Nur77-GFP^low^ and Ly49H^-^ NK cells. **C)** Spearman correlations of early antigen primed NK cells with *in vitro* stimulated NK cells using RNA-seq (Top) or ATAC-seq (Bottom). **D)** Representative example DA regions (*Irf4*, *Crtam*) with quantification of peak signals. **E)** Scatterplot of ATAC-seq from Ly49H+ Nur77-GFPhigh versus Ly49H+ Nur77-GFPlow NK cells and unstimulated versus anti-NK1.1 stimulated NK cells. Colored circles are significant (padj <0.05), and highlighted circles show regions with differential STAT4 binding in Fig. 2A. **F)** Scatterplot of RNAseq and ATACseq from NK cells stimulated with aNK1.1 or aLy49H. Colored circles represent significant genes/regions (padj <0.05).

**Supp. Fig. 4:**
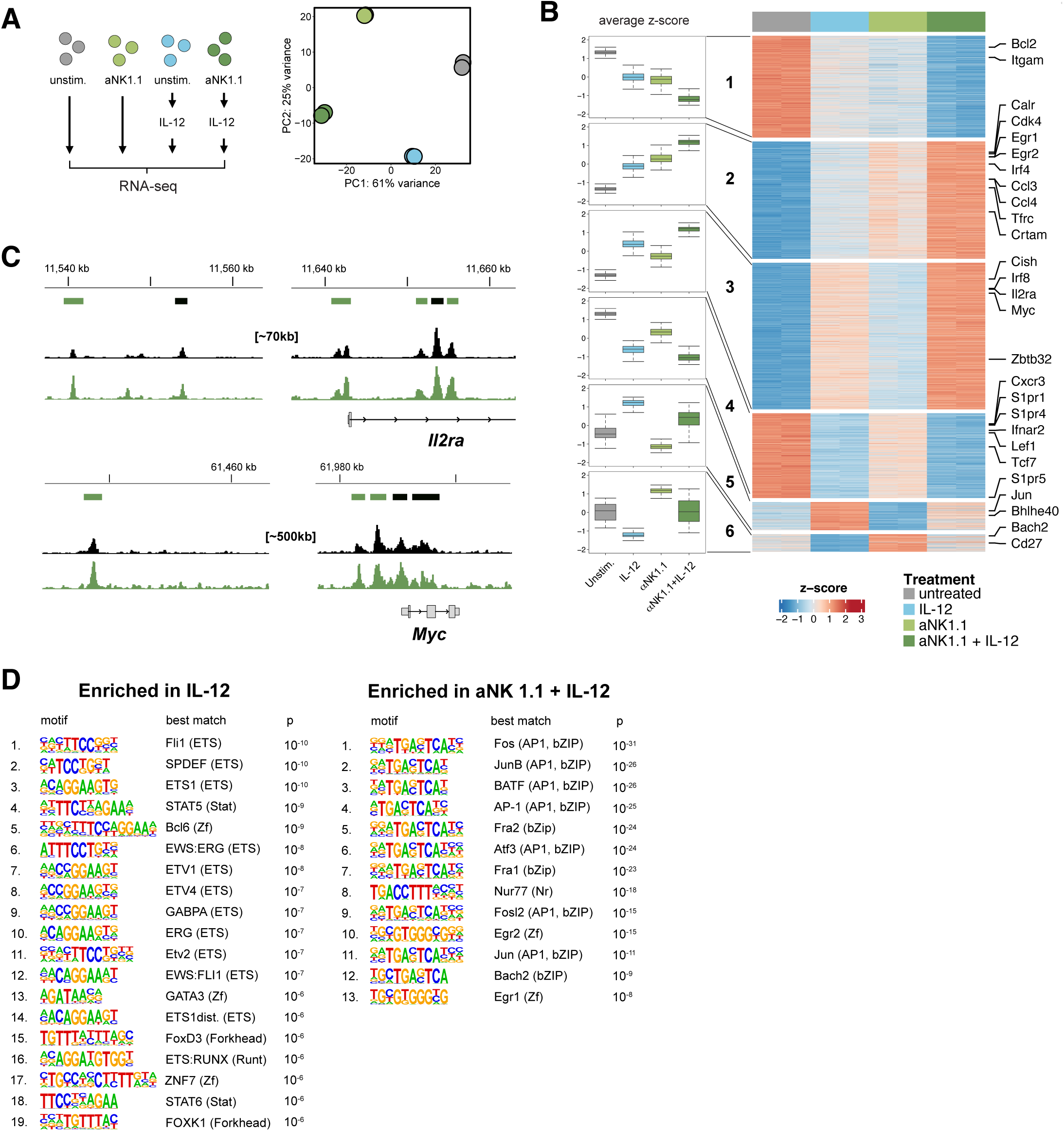
**A)** Schematic and PCA of sequential RNAseq experiment with aNK1.1 and IL-12 stimulation. **B)** Heatmap of RNAseq from sequential stimulation. Grouping depending on different expression patterns. **C)** Representative peaks of *Il2ra* and *Myc* locus. Green bars represent AP-1 sites. **D)** Table of HOMER known motif analysis. The top 20% peaks denoted above were analyzed for enrichment for known motifs in HOMER database, with the other STAT4 binding atlas peaks as background. The top significant known motifs are shown.

**Supp. Fig. 5:**
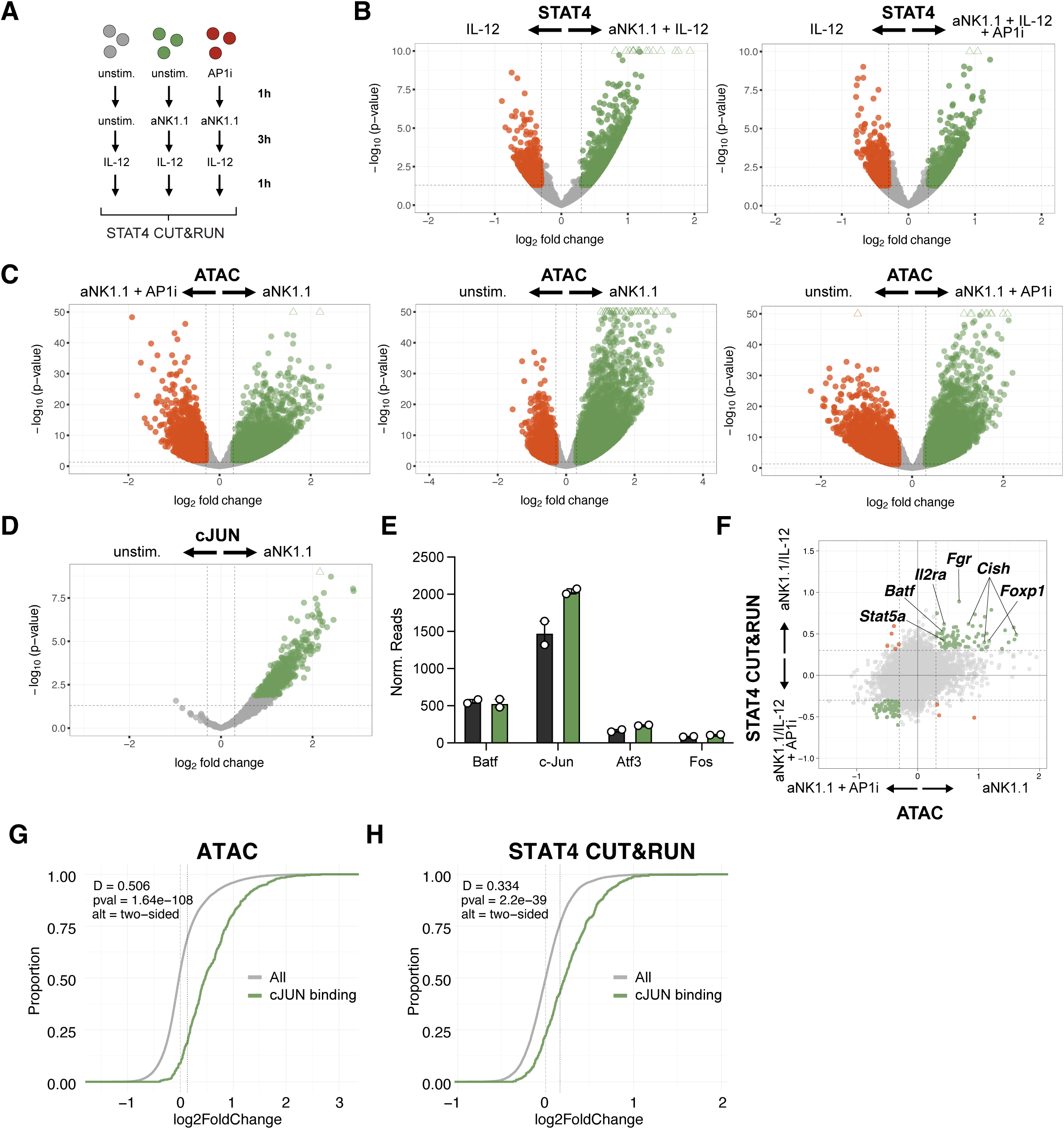
**A)** Schematic of sequential STAT4 CUT&RUN experiment with AP1i, aNK1.1 and IL-12 stimulation. **B)** Volcanoplot of STAT4 CUT&RUN in different conditions. Colored circles are p < 0.05 and log2foldchange > 0.3. **C)** Volcanoplot of ATACseq in different conditions. Colored circles are p < 0.05 and log2foldchange > 0.3. **D)** Volcanoplot of cJUN CUT&RUN in unstimulated or aNK1.1 stimulated NK cells. **E)** Normalized Reads of AP-1 transcription factor genes in unstimulated NK cells or NK cells stimulated with aNK1.1 (Experiment from Fig. 2D). **F)** Scatterplot of STAT4 CUT&RUN and ATACseq for NK cells stimulated with aNK1.1 + IL-12 vs. aNK1.1 + IL-12 + AP1i and aNK1.1 vs. aNK1.1 + AP1i, respectively. **G)** ECDF plot of ATACseq log2foldchange of unstim. vs. aNK1.1 stimulated NK cells subsetted for peaks bound by cJUN after antigen stimulation (green) vs. all peaks (grey). Shift indicates that peaks bound by cJUN gain accessibility. **H)** ECDF plot of STAT4 CUT&RUN log2foldchange of IL-12 vs. aNK1.1 + IL-12 stimulated NK cells subsetted for peaks bound by cJUN after antigen stimulation (green) vs. all peaks (grey). Shift indicates that peaks bound by cJUN show increased STAT4 genomic binding. Statistics for ECDF plots are derived from Kolmogorov-Smirnov test.

**Supp. Fig. 6:**
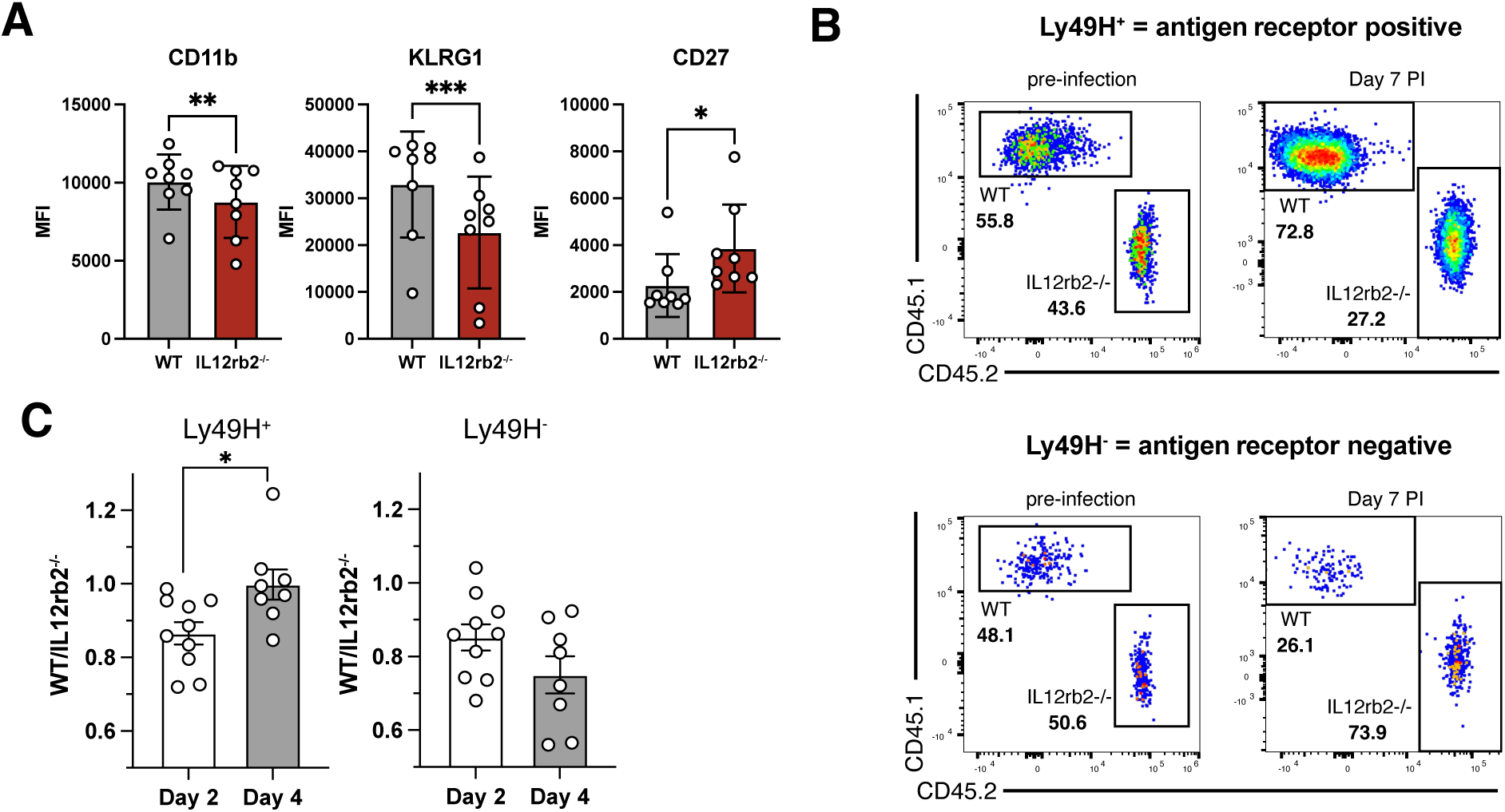
**A)** Expression of key maturation markers on day 7 post infection for Fig. 3A. **B)** Representative flow plots for Fig. 3B). **C)** WT/Il12rb2^-/-^ ratios in Ly49H^+^ and Ly49H^-^ NK cells on day 4 post infection in WT: IL12rb2^-/-^ chimera. Experiments in Fig. S4A) and Fig. S4C) is pooled from two independent experiments. Error bars represent SEM. Significances are calculated using paired T-test (same biological replicates). **** p < 0.0001, *** p < 0.001, ** p < 0.01, * p < 0.05.

**Supp. Fig. 7:**
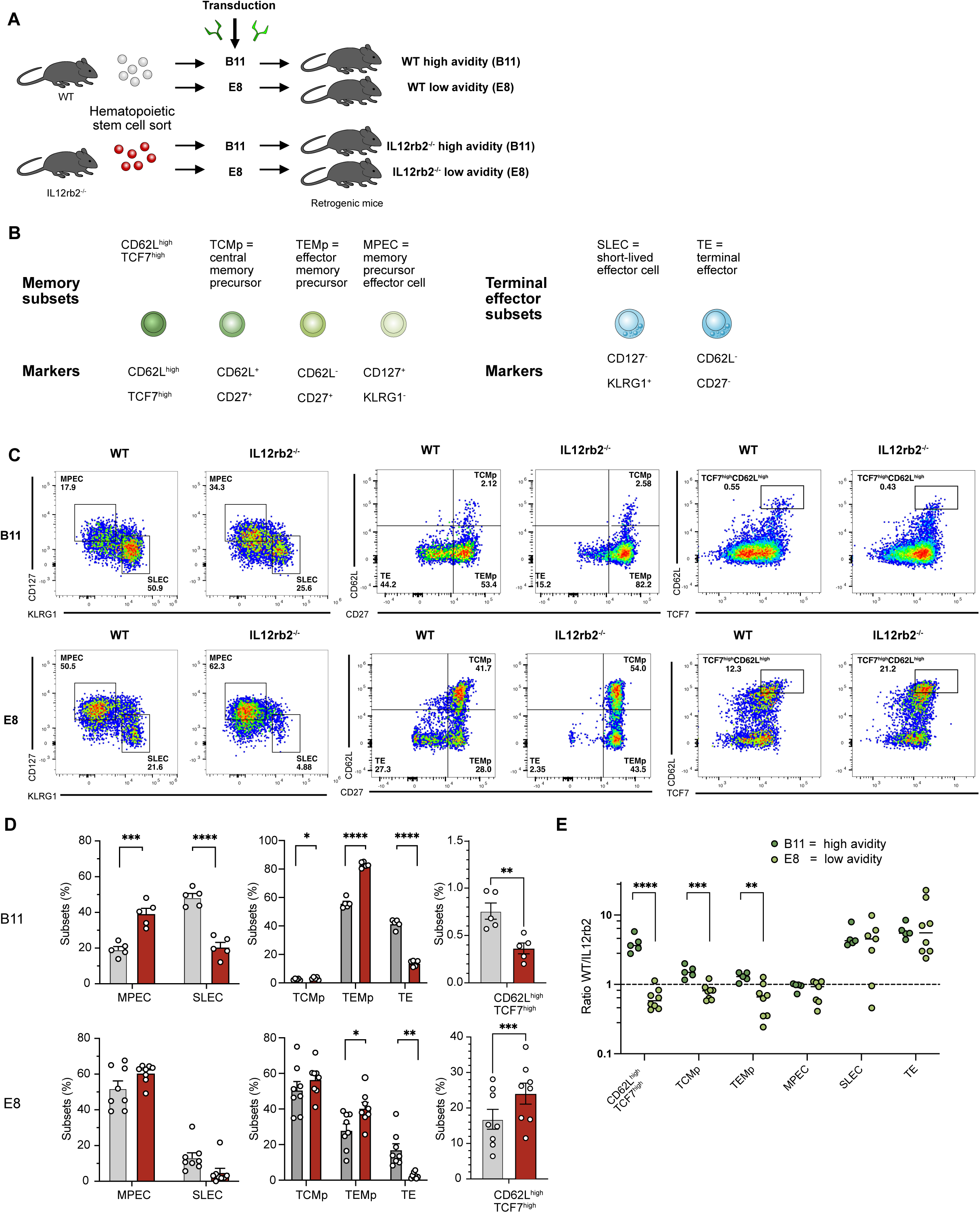
**A)** Schematic of retrogenic mouse generation from WT and *Il12rb2^-/-^*bone marrow. **B)** Schematic of different memory precursor and terminal effector subsets. We use and compare three different classification used in literature: classification 1) using markers CD62L and TCF7: CD62L^high^ TCF7^high32,33^; classification 2) using markers CD62L and CD27: TCMp, TEMp and TE^33,64^; classification 3) using markers CD127 and KLRG1^6^. **C)** Representative flow plots. All cells from all mice from one experiment were concatenated and dot numbers were normalized to 3500 for all conditions but CD62L^high^TCF7^high^ (10.000 dots for B11 for better visual representation). **D)** Average phenotype of high-avidity (B11) and low-avidity (E8) WT and *Il12rb2-/-* CD8^+^ T cells using MPEC/SLEC, TCMp/TEMp/TE or CD62L^high^TCF7^high^ classification. **E)** Ratios (WT/*Il12rb2^-/-^*) of absolute subset output for different subsets. Data in Fig. S5D) and S5E) are representative of two independent experiments. Error bars represent SEM. Significances are calculated using unpaired T-test. **** p < 0.0001, *** p < 0.001, ** p < 0.01, * p < 0.05.

**Supp. Fig. 8:**
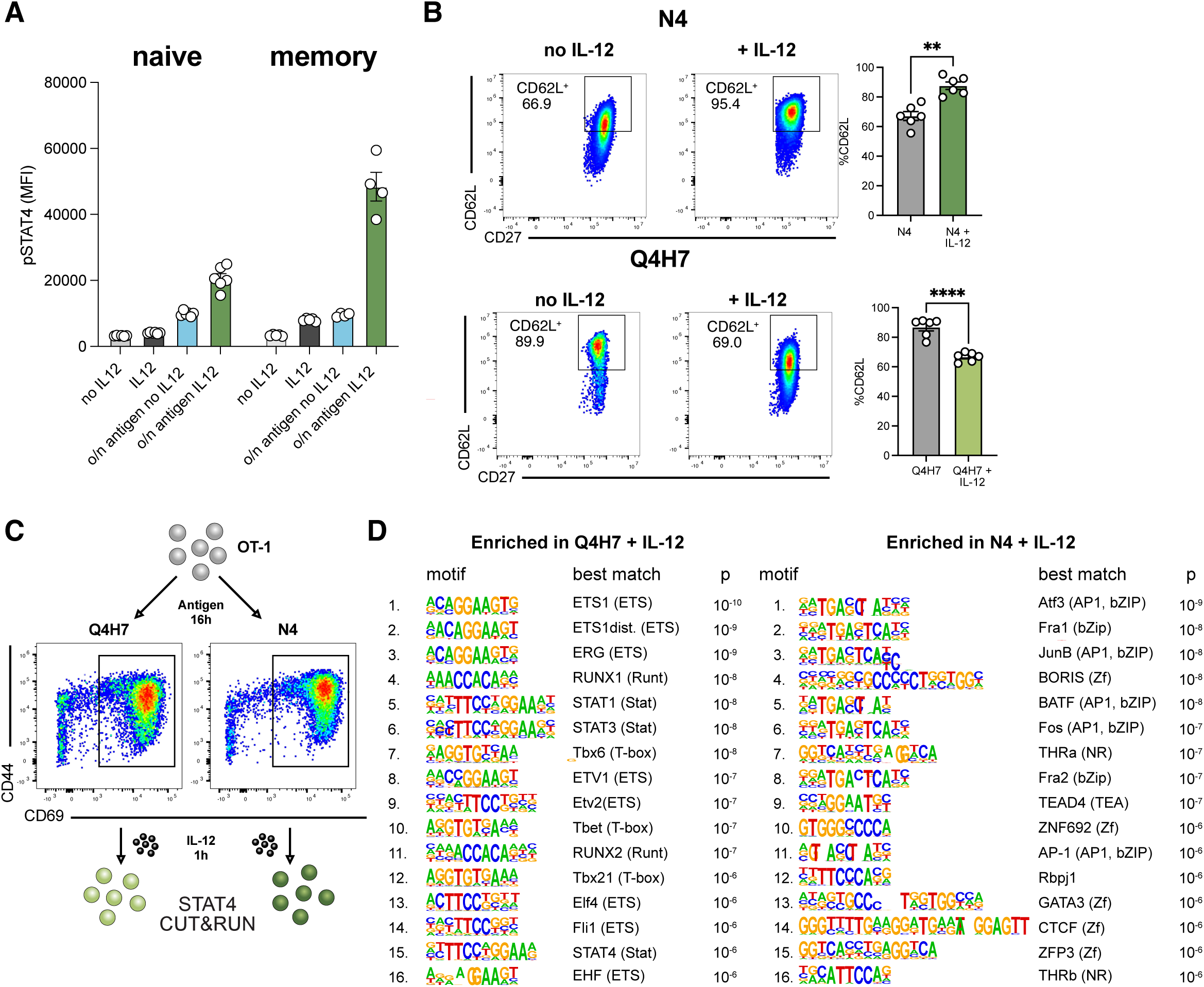
**A)** Naïve OT-1 or memory OT-1 (>d35 post infection with L.m.-SIINFEKL) were directly stimulated with IL-12 or incubated with SIINFEKL peptide overnight before IL-12 stimulation and stained for pSTAT4. **B)** Representative flow plots for CD62L expression for Fig. 4D. **C)** Schematic and representative flow plots of high- and low-avidity stimulated OT-1 T cells at time of STAT4 CUT&RUN. **D)** Table of HOMER known motif analysis. The top 20% peaks denoted in Figure 4E were analyzed for enrichment for known motifs in HOMER database, with the other STAT4 binding atlas peaks as background. The top significant known motifs are shown. Data in Fig. S8A) is representative of three and S8B) of two experiments. Error bars represent SEM. Significances were calculated using paired T-test (same biological replicates). **** p < 0.0001, *** p < 0.001, ** p < 0.01, * p < 0.05.

